# Decoding neuronal diversity by single-cell Convert-seq

**DOI:** 10.1101/600239

**Authors:** Joachim Luginbühl, Tsukasa Kouno, Rei Nakano, Thomas E Chater, Divya M Sivaraman, Mami Kishima, Filip Roudnicky, Piero Carninci, Charles Plessy, Jay W Shin

## Abstract

The conversion of cell fates is controlled by hierarchical gene regulatory networks (GRNs) that induce remarkable alterations in cellular and transcriptome states. The identification of key regulators within these networks from myriad of candidate genes, however, poses a major research challenge. Here we present Convert-seq, combining single-cell RNA sequencing (scRNA-seq) and pooled ectopic gene expression with a new strategy to discriminate sequencing reads derived from exogenous and endogenous transcripts. We demonstrate Convert-seq by associating hundreds of single cells during reprogramming of human fibroblasts to induced neurons (iN) with exogenous and endogenous transcriptional signatures. Convert-seq identified GRNs modulating the emergence of developmental trajectories and predicted combinatorial activation of exogenous transcription factors controlling iN subtype specification. Functional validation of iN subtypes generated by novel combinations of exogenous transcription factors establish Convert-seq as a broadly applicable workflow to rapidly identify key transcription factors and GRNs orchestrating the reprogramming of virtually any cell type.

## Introduction

Fully differentiated cells can be reprogrammed to alternative fates via ectopic expression of defined combinations of transcription factors and/or microRNAs (Chanda et al., 2014; Hu et al., 2015; Mertens et al., 2015; Pang et al., 2011; Pfisterer et al., 2011; Ryoji Amamoto and Arlotta, 2014; Victor et al., 2014; Vierbuchen et al., 2010; Xue et al., 2013). Reprogrammed cells offer invaluable alternative routes for the treatment of various diseases and represent a substantial improvement to the field of disease modeling (Masserdotti et al., 2016). However, despite the establishment of new in silico prediction tools (Rackham et al., 2016), it has remained a challenge to identify essential genetic drivers that determine specific subtypes of cells during reprogramming.

Cell reprogramming is of particular interest for human neurons, which are notoriously difficult to obtain from patients and cannot be expanded in culture (Ryoji Amamoto and Arlotta, 2014). Since the first report on reprogramming of induced neurons (iN), several studies have succeeded in identifying defined combinations of neurogenic transcription factors that allow the reprogramming of specific neuronal subtypes (Masserdotti et al., 2016). For instance, distinct sets of neurogenic transcription factors have been shown to create dopaminergic neurons and cholinergic motor neurons that are selective targets of degeneration in patients with Parkinson’s disease and amyotrophic lateral sclerosis (ALS), respectively (Caiazzo et al., 2011; Jiang et al., 2015; Kim et al., 2011; Liu et al., 2013, 2016; Mazzoni et al., 2013; Pfisterer et al., 2011; Son et al., 2011). However, an inherent limitation of studies aiming to identify key transcription factors controlling cell reprogramming is that the identification of candidates from myriad of transcription factors still is based on trial and error. This is exacerbated by the time and labor intensive retesting and validation of newly identified candidates. Pooled screens, instead of one-by-one tests, offer a more efficient and scalable strategy, but they either rely on simple phenotypic readouts or require specific markers to enrich for target cell populations, which are often not available (Chen et al., 2015; Hsu et al., 2014; Liu et al., 2018; Shalem et al., 2015). This is particularly problematic with regard to reprogramming, which is intrinsically inefficient, resulting in heterogenous cell populations containing unreprogrammed, partially reprogrammed, and successfully reprogrammed cells.

To address this challenge, we developed Convert-seq, combining ectopic expression of transcription factors with single-cell RNA sequencing (scRNA-seq) to retrospectively associate single cells with their exogenous and endogenous expression profiles. Convert-seq represents a pooled (multiplexed) screening strategy that allows to identify transcription factors and GRNs governing the reprogramming of many different celltypes in a single experiment. We demonstrate Convert-seq in iN generated by the pooled infection of human fibroblasts with 20 pro-neuronal transcription factors using two single-cell sequencing platforms, the Fluidigm C1 system based on microfluidic technology and the 10× Genomics system based on nano-droplets. By association of different transcriptional states and developmental trajectories emerging during reprogramming with distinct exogenous and endogenous transcriptional signatures, we identified defined combinations of key transcription factors orchestrating the reprogramming of human fibroblasts into multiple neuronal subtypes. Our data demonstrate the ability of Convert-seq to efficiently gain rich insight into biological processes governing cell plasticity and cell fate decisions and to systematically dissect GRNs controlling subtype specification during cell reprogramming.

## Results

### Generation of a heterogenous population of human induced neurons

To enable systematic identification of combinations of transcription factors governing the reprogramming of human fibroblasts into specific neuronal subtypes, we generated a pool of lentiviruses (hereafter termed transcription factor-pool or TF-pool) encoding 20 pro-neuronal transcription factors. We included the pioneer factor *ASCL1* (Wapinski et al., 2013) and several transcription factors that increase the efficiency of neuronal reprogramming (*POU3F2, ZIC1, OLIG2, NEUROD1*) (Pang et al., 2011; Vierbuchen et al., 2010). To promote reprogramming of different subtypes of neurons, we further included transcription factors which have been previously shown to convert human fibroblasts into GABAergic (*DLX1*, *DLX2*) (Victor et al., 2014), cholinergic (*NEUROG2*, *ISL1*) (Liu et al., 2013; Son et al., 2011), serotonergic (*FEV*) (Xu et al., 2016) or dopaminergic (*FOXA2*, *NR4A2*, *PITX3*) (Liu et al., 2012; Pfisterer et al., 2011) neurons when co-expressed with other transcription factors. Additionally, we selected 7 transcription factors which have not yet been used to generate specific neuronal subtypes, but showed significant up-regulation during the differentiation of human induced pluripotent stem cells (iPSC) into neuronal progenitor cells (NPC) (Figure S1A). Each transcription factor was expressed under the elongation factor 1 alpha (EF1A) promoter together with a gene encoding Venus yellow fluorescent protein (YFP) (Figure S1B). Based on quantifications of YFP expression, we tuned the multiplicity of infection (MOI) of each lentivirus to allow ~85% of fibroblasts to express mixed combinations of 2 - 6 transcription factors after stochastic infection with the TF-pool (Figure 1A and Figure S1C). Using successive application of neuronal induction medium for 14 days followed by neuronal maturation medium for 7 days, we successfully transformed fibroblasts into cells exhibiting hetergeneous neuronal morphologies and expressing canonical neuronal marker genes (Figures 1B). At 9 days post-infection (dpi), 78.6% of fibroblasts (hereafter termed transcription factor-induced cells or TFi) stained positive for the immature neuronal marker TUBB3 and 34.7% expressed MAP2, a microtubule-associated protein expressed specifically in neurons (Jeff et al., 1988) (Figure 1C). By 21 dpi, TUBB3+ and MAP2+ ratios increased to 93.5% and 42.4%, respectively, indicating progressive differentiation towards the neuronal lineage. Consistent with the adoption of a neuronal fate, quantitative polymerase chain reaction (qPCR) revealed gradual upregulation of pan-neuronal marker genes (*MAP2*, *NRCAM*, *NEUN*, *SYN1*) and established neuronal subtype marker genes (*SLC17A7* [glutamatergic neurons], *GABRA1* [GABAergic neurons], *TH* [dopaminergic neurons] and *CHAT* [cholinergic neurons]) as well as gradual downregulation of fibroblast marker gene expression (*VIM, SNAI1*) in TFi starting at 7 dpi (Figures 1D).

**Figure 1.**
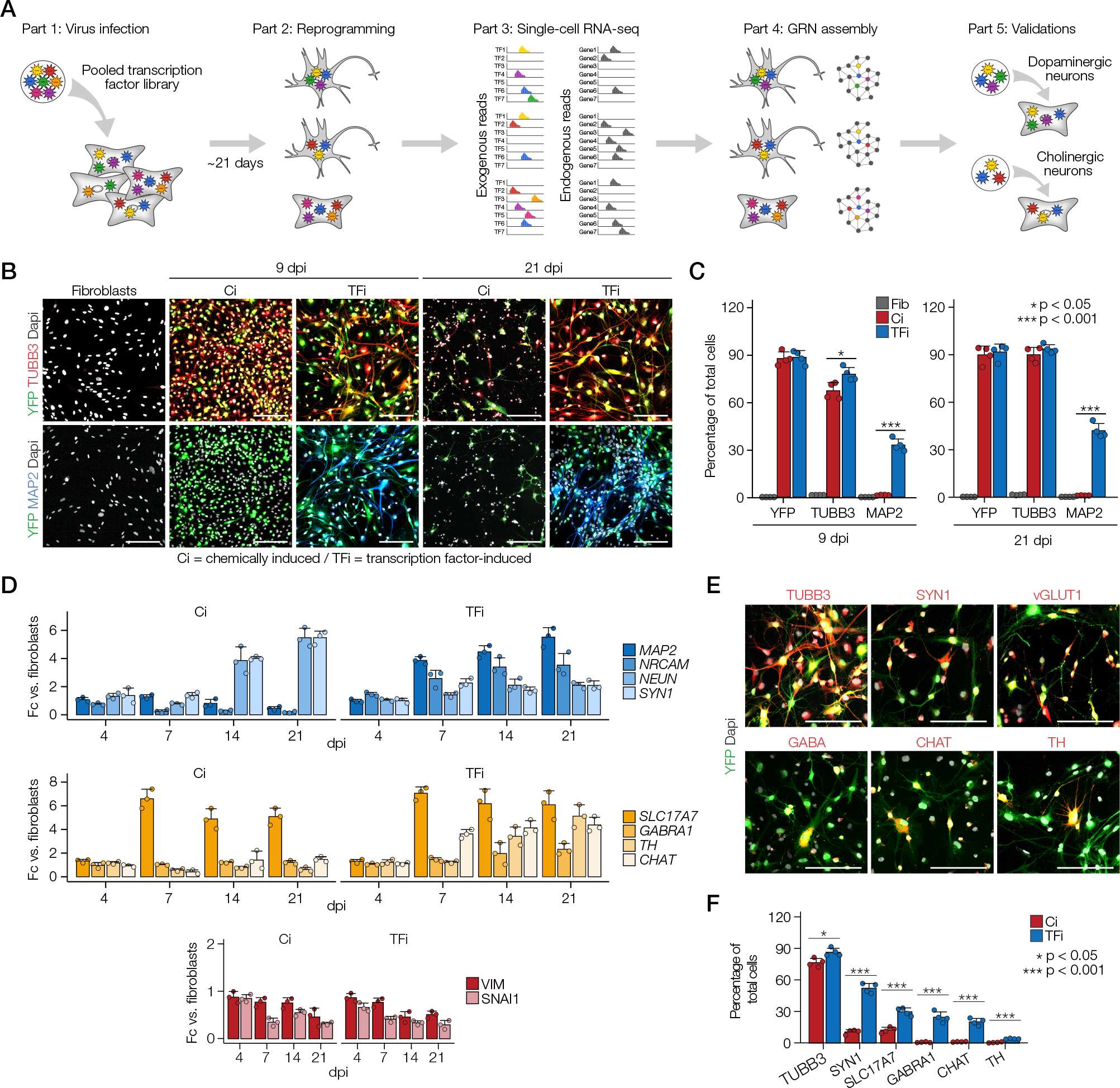
Generation of a heterogenous population of human induced neurons. (A) Overview of Convert-seq. (B) Immunostaining for canonical neuronal marker genes of fibroblasts at day 0 and Ci and TFi at 9 dpi and 21 dpi. Scale bars, 100μm. YFP (green) marks infected cells and cell nuclei were visualized using DAPI nuclear stain (grey). (C) Quantification of immunostainings in B. *n* = 4 independent experiments, error bars represent mean + SD. (D) qPCR for pan-neuronal marker genes (*MAP2*, *NRCAM*, *NEUN*, *SYN1*; top), canonical neuronal subtype markers (*SLC17A7*, *GABRA1*, *TH*, *CHAT*; middle) and canonical fibroblast markers (*VIM*, *SNAI1*; bottom). n = 3 independent experiments, error bars represent mean + SD. (E) Immunostainings for canonical neuronal subtype markers (red) of TFi at 21 dpi. Scale bars, 100μm. YFP (green) marks infected cells and cell nuclei were visualized using DAPI nuclear stain (grey). (F) Quantification of immunostainings in E. Error bars represent mean + SD.

Previous studies have shown that mouse and human fibroblasts can be directly converted to induced neurons (iN) solely by chemical cocktails of small molecules (Hu et al., 2015; Li et al., 2015). Thus, to discriminate between transcription factor-mediated and chemical-mediated conversion, we separately cultured fibroblasts transduced with lentiviruses expressing YFP only (hereafter termed chemically induced cells or Ci) using the same neuronal induction regime. Interestingly, we found that similar to TFi, application of small molecules and growth factors alone induced the development of TUBB3+ neurites, upregulated the expression of *NEUN*, *SYN1*, *SLC17A7* and *TUBB3* and downregulated the expression of *VIM* and *SNAI1* in Ci (Figures 1B-1D). However, Ci failed to express *MAP2* at any time-point analyzed, indicating that the small molecules we used were insufficient to elicit neuronal maturation on their own. Consistently, neuronal complexity, measured as number of branch points and average neurite length, was significantly increased at 9 dpi (8.5-fold and 1.8-fold, respectively) and at 21 dpi (5.6-fold and 4.8-fold, respectively) when fibroblasts were infected with the TF-pool compared to application of small molecules alone (Figures S1E-S1F).

### Molecular characterization of induced neurons using scRNA-seq

Immunofluorescence for canonical neuronal subtype markers at 21 dpi revealed that TFi differentiated into a heterogeneous population of cells expressing glutamatergic (vGLUT1+; ~29%), GABAergic (GABA+; ~23%), cholinergic (CHAT+; ~19%) and dopaminergic (TH+; ~4%) subtype-specific genes (Figures 1E-1F). In contrast, only ~15% of Ci expressed vGLUT1 at 21 dpi and none of the other subtype-specific genes was induced by small molecules.

To gain further insight into the molecular heterogeneity of TFi and Ci, we used droplet-based massively parallel scRNA-seq (Zheng et al., 2017) to profile 2,092 Ci and 1,900 TFi at 14 dpi (Table S1). Batch-corrected clustering of all cells that passed quality control using *t*-Distributed Stochastic Neighbor embedding (*t-*SNE) separated Ci and TFi along *t-*SNE 2 (Figures 2A and S2A-D). Correlation analysis of top genes along *t-*SNE 2 (Pearson correlation; p < 10^−7^) revealed activation of genes relevant for neuronal differentiation (*NRCAM, STMN2, SST, DKK3*) and synapse formation (*SYT1*, *SERPINI1, SYNGR1, SYNPO2*), which was accompanied by a decrease in fibroblast-specific genes (*SNAI2, THY1*) (Figure 2B). Other suppressed genes included extracellular matrix genes (*COL3A1, COL15A1, EDIL3, HAPLN1*, *CPXM1* and *MFAP4)*, reflective of the morphological changes that occur during the conversion of iN. t-SNE further partitioned TFi cells into one main cluster (CL3) and several transcriptionally distinct clusters (CL4-CL9). CL5 dominantly expressed cell cycle-related genes including *MIKI67* and *TOP2A* and showed enrichment of Gene ontology (GO) terms related to cell division, indicating that these cells – possibly un-reprogrammed fibroblasts – remained in a mitotic stage (Figures 2C-D; Table S2). Importantly, consistent with morphology and marker expression stainings, clusters CL4 and CL6 – CL9 all showed enrichment in terms related to nervous system development and neurogenesis. To annotate clusters CL4 – CL9, we interrogated top differentially expressed genes (Seurat; p < 10^−20^) for known neuronal subtype marker genes. This revealed CL4 to associate with immature neurons, CL6 with the motor neuron program (*CHRNA5*, *CP*), CL7 with the GABAergic neuron program (*ANK3*, *NKAIN4*), CL8 with the glutamatergic neuron program (*SLC6A17*, *INSM1*) and CL9 with the dopaminergic neuron program (*DRD4, SEMA3G*) (Figures 2E-F and S2E).

**Figure 2.**
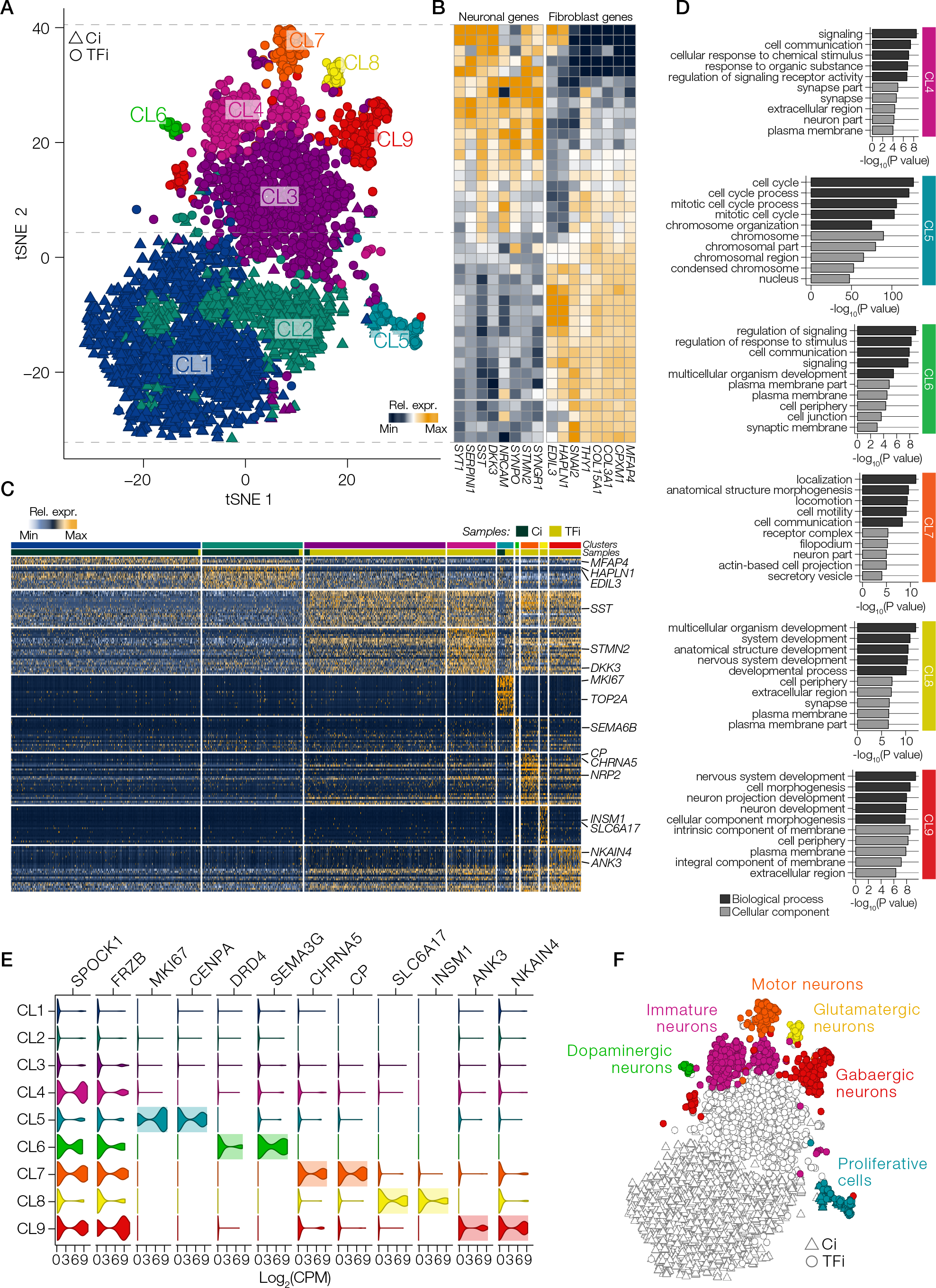
Molecular characterization of induced neurons using scRNA-seq. (A) Visualization of droplet-based scRNA-seq data from Ci and TFi at 14 dpi using t-SNE (*n* = 3865 cells). The detected clusters are indicated by different colors. (B) Heat map of the relative expression of canonical fibroblast and neuron markers along t-SNE 2. (C) Heat map of the relative expression of top marker genes for each cluster in the t-SNE plot in (A). (D) GO analysis of cluster-specific marker genes in clusters CL4 – CL9. Shown are top 5 GO terms related to biological process (dark grey) and cellular component (light grey) for each cluster; colors as in (A). (E) Violin plots of log_2_−transformed CPM values of marker genes in all clusters. (F) Annotation of TFi clusters CL4 – CL9 based on genes differentially expressed between each cluster.

Together, these data revealed that transduction of our TF-pool converted fibroblasts into a population of cells exhibiting a high degree of heterogeneity and distinct neuronal subtype-specific molecular signatures.

### Convert-seq Detection of Exogenous and Endogenous Transcripts

To decode the identity of exogenous transcription factors in individual cells, we designed Convert-seq, combining the stochastic nature of pooled, virus-based screens with massively parallel scRNA-seq (Figure 1A). Based on full-length reads, we extracted nucleotides at the 5‘ and 3‘ junctions of exogenous ORFs and assembled artificial transcript models of endogenous and exogenous reference sequences (Figure 3A). Alignment of reads to these specific 5’ and 3’ junctions allows to discriminate between exogenous and endogenous gene expression. To benchmark the accuracy and sensitivity of Convert-seq, we profiled gene expression of bulk fibroblasts infected with single transcription factors, two combinations of 10 transcription factors, and the complete TF-pool at two multiplicity of infections (MOI) (Figures 3B and S3A). We also included chemically-induced (Ci) cells and uninfected fibroblasts as controls. Sequence alignment to the artificial transcript model using Bowtie (Langmead et al., 2009) revealed specific detection of exogenous transcription factors, but resulted in 24.4% false-negative events, possibly due to the exclusion of multi-mapping reads (Figure 3C). To reduce the number of false-negative events, we implemented the pseudo-alignment tool Kallisto, providing the advantage of assigning multi-mapping reads to transcripts without pinpointing exactly how the sequences of the reads and transcripts align (Bray et al., 2016). Pseudoalignment with Kallisto reduced the occurrence of false negative events to 6.3% and yielded highly specific and sensitive detection efficiencies of 97.5% and 87.5% in individual infection and pooled infection samples, respectively (Figures 3D and S3B).

**Figure 3.**
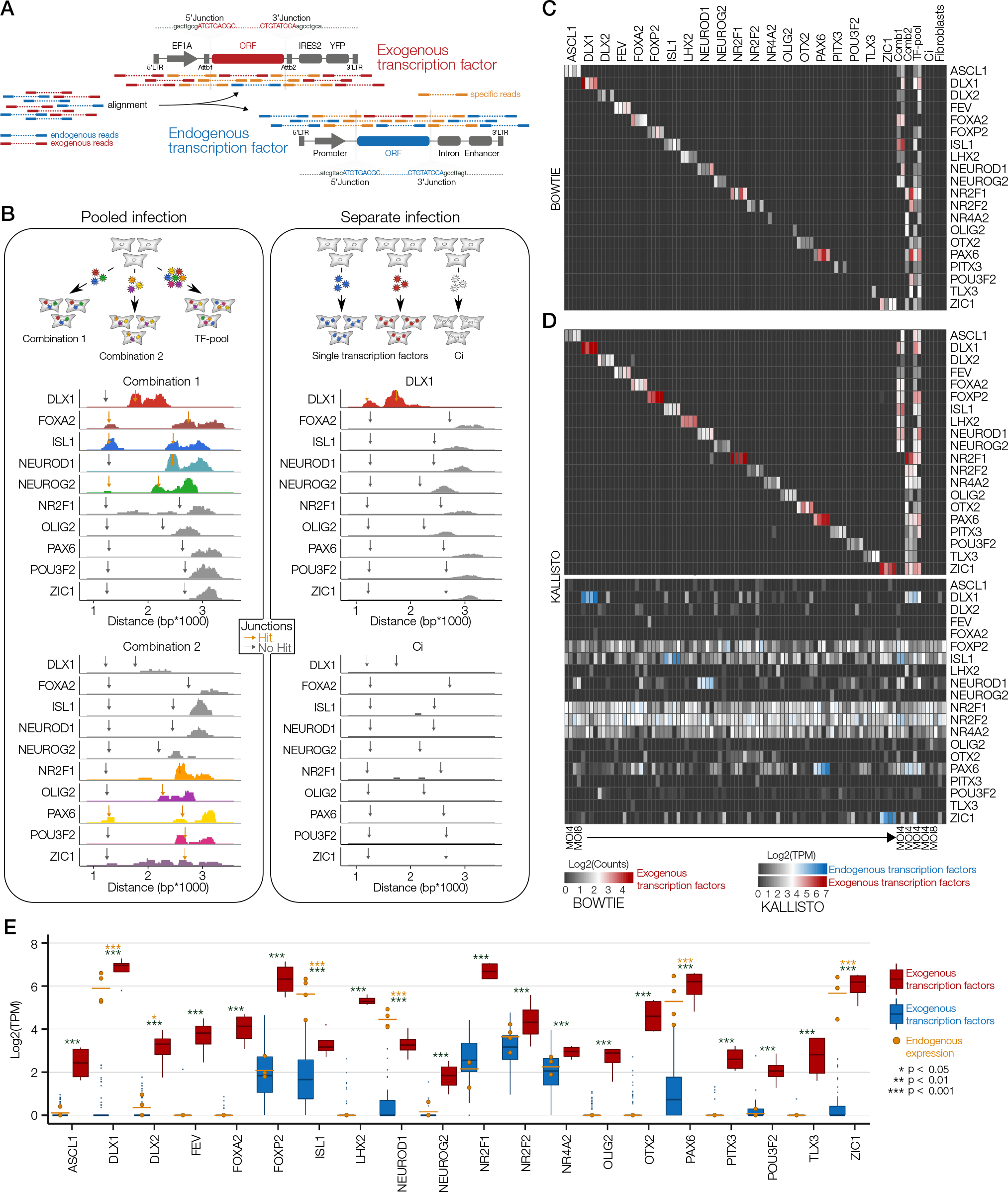
Convert-seq Detection of Exogenous and Endogenous Transcripts. (A) Schematic depicting the strategy to distinguish exogenous and endogenous sequencing reads. (B) Bulk Convert-seq on pooled and separately infected fibroblasts. Horizontal dimension; distance from the 5‘ end of the EF1A promoter, vertical dimension; number of aligned paired-end reads. Gray arrows (no overlap) and golden arrows (overlap) mark 5’ and 3’ junctions of exogenous ORFs. (C-D) Heat maps showing log_2_−transformed count values of exogenous transcription factors after alignment using Bowtie (C) and log_2_−transformed TPM values of exogenous and endogenous transcription factor pairs after trimming junction sequences to ~100 base pairs and alignment using Kallisto (D). For individually infected fibroblasts and Ci, 2 replicates at an MOI of 4 and 2 replicates at an MOI of 8 were included. For pooled infected fibroblasts, 2 replicates at an MOI of 4 were included. (E) Box plots showing increased exogenous (red) versus endogenous (blue) expression of all transcription factors across all individually infected fibroblasts. Golden dots show endogenous expression in samples infected with the corresponding exogenous transcription factors. Error bars represent mean + SD.

To further confirm specific detection of lentivirus-mediated expression, we revealed significantly higher mean expression of exogenous transcription factors compared to the mean expression of corresponding endogenous transcription factors (p < 0.05; Figure 3E). Interestingly, additional infection experiments revealed that for several genes, ectopic expression failed to activate endogenous expression, while we observed upregulation of endogenous *DLX1*, *DLX2*, *ISL1*, *NEUROD1*, *PAX6* and *ZIC1* transcripts after infection with their exogenous counterparts (Figure S3C).

### Induced neurons resemble primary human newborn neurons in global gene expression

Convert-seq requires full-length coverage of transcripts to obtain sequencing reads falling into specific 5’ and 3’ junction sequences. Accordingly, detection of exogenous genes from the 3’-biased droplet-based scRNA-seq data was inefficient (Figure S3D). Therefore, we sequenced transcription factor-induced cells (TFi) and chemically induced cells (Ci) at an early (9 dpi) and late (21 dpi) time point (216 cells and 152 cells, respectively) on the Fluidigm C1 platform using Smartseq2 technology (Figure 4A). Unsupervised clustering of all cells that passed quality control using *t-*SNE identified 5 transcriptionally distinct clusters of cells that separated largely by type rather than batch (Figures 4B and S4A-F). GO analysis of differentially expressed genes between all clusters (fold-change > 2; p < 10^−4^) revealed that CL1 dominant in unreprogrammed fibroblasts specifically expressed genes related to cell cycle and cell division, whereas CL2 dominant in Ci at 9 dpi was characterized by specific expression of genes related to general developmental processes without bias towards nervous system development (Figures 4C and S4G-H; Table S3). CL3 dominant in TFi at 9 dpi expressed genes related to the inflammatory response and cell cycle regulation, indicating inefficient initiation of the reprogramming process. Similar to CL2, CL4 dominant in TFi at 21 dpi expressed genes related to general developmental processes. Finally, CL5 expressed genes related to neuron development and neuron maturation, suggesting that CL5 contained cells that were most efficiently reprogrammed towards the neuronal lineage.

**Figure 4.**
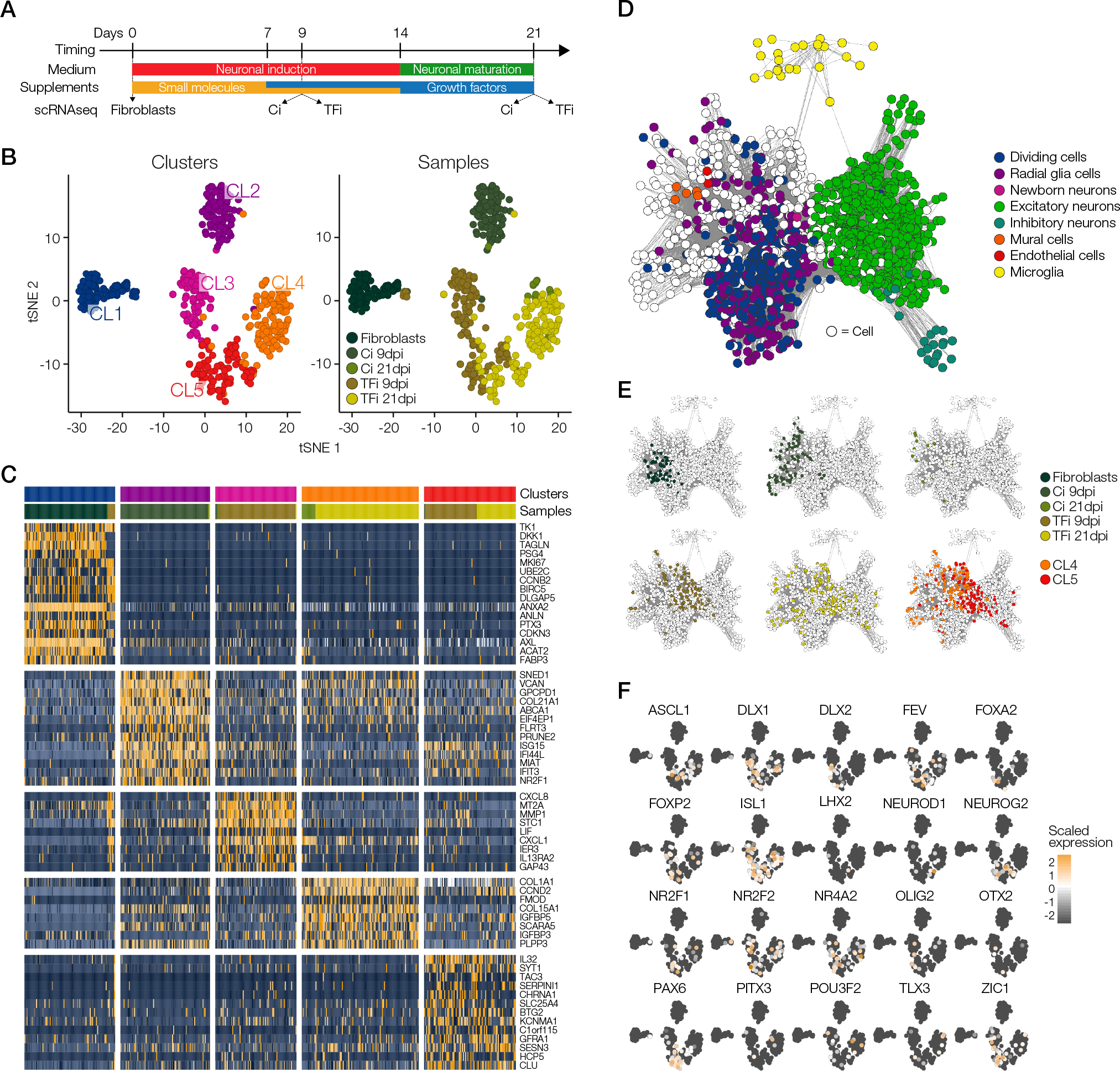
Induced neurons resemble primary human newborn neurons in global gene expression. (A) Diagram of the differentiation protocol of TFi and Ci, depicting the samples used for the time-course scRNAseq experiment. (B) t-SNE 2D cell maps of the time-course data. Left: cells were colored by cluster identity. Right: cells were colored by sample identity. Fibroblasts (n = 78 cells), Ci at 9 dpi (n = 87 cells), TFi at 9 dpi (n = 129), Ci at 21 dpi (n = 15 cells), TFi at 21 dpi (n = 137 cells). (C) Heat map of the relative expression of top marker genes for each cluster in the t-SNE plot in (B). (D) Lineage relationships of fibroblasts, Ci, TFi and primary cortical and medial ganglionic eminence cells (*n* = 4261 cells) in an adjacency network. (E) Visualization of relative expression values of exogenous transcription factors on *t*-SNE plots.

As an alternative approach to assess reprogramming efficiency, we used a publicly available scRNA-seq dataset of human primary cortical and medial ganglionic eminence samples to infer lineage relationships among the cells in an adjacency network on the basis of pairwise correlations between cells (Nowakowski et al., 2017) (Figure 4D). This analysis revealed that while Ci correlated with alternative cell types including endothelial cells and mural cells, TFi showed progressive correlation with newborn neurons, excitatory neurons and inhibitory neurons. Consistent with our previous results (Figure S4G-H), we found that TFi exhibiting transcriptional profiles related to cell cycle regulation and inflammatory response showed stronger correlation with neuronal progenitors and newborn neurons, whereas the majority of TFi exhibiting transcriptional profiles related to neuron development and neuron maturation correlated with excitatory and inhibitory neurons (Figure 4E).

### Association of developmental trajectories with exogenous expression profiles

A previous study in mouse embryonic fibroblasts showed that reprogramming to neurons occurs through a continuum of intermediate states, whereby ectopic expression of specific exogenous transcription factors is required to suppress aberrant developmental trajectories and maintain neuronal identity (Treutlein et al., 2016). To analyze intermediate states in our time-course data and simultaneously associate the expression of exogenous transcription factors with distinct developmental trajectories during reprogramming, we first used single-cell Convert-seq to assign individual cells with exogenous transcription factors and corresponding endogenous gene expression profiles (Figure 4F). Next, we used the Monocle algorithm (Trapnell et al., 2014) to place the cells in pseudo-temporal order based on differentially expressed genes between 9 dpi and 21 dpi (qval < 0.01) (Figure S5A). This revealed that cells bifurcated into two divergent developmental trajectories shortly after initiation of reprogramming. Using branch-specific differential gene expression (fold-change > 4; p < 10^−2^), we found that branch 1 cells contained mostly TFi and Ci from 9 dpi and expressed genes involved in viral expression, ribosome biogenesis and cell cycle control, whereas branch 2 cells contained TFi and Ci from both 9 dpi and 21 dpi, and expressed genes related to nervous system development (Figures S5B-C; Table S4). Next, we correlated differentially expressed genes across pseudotime. Unexpectedly, positively correlating genes (r > 0.2) were enriched for GO terms related to various developmental processes (Figures S5D-E, Table S5), implying the emergence of distinct developmental trajectories during reprogramming. Indeed, assessment of gene expression levels along pseudotime confirmed that neurogenic genes (*NRCAM*, *SFRP1*, *SNAP25* and *SYT1*) and genes regulating the development of mesodermal tissues including bone (*BMP4*), kidney (*FAT4*) and endothelial cells (*PGF* and *VEGFA*) were consistently up-regulated and followed a similar expression trajectory pattern during the reprogramming process (Figure S5F-H).

To identify exogenous transcription factors governing the emergence of distinct developmental trajectories, we reordered the cells using a set of 3,322 genes previously implicated in various developmental processes (Figure 5A; Table S6). This revealed bifurcation of cells into two main trajectories, separating the majority of Ci and part of early TFi (branch 1) from the rest of TFi (branch 2). Branch-specific GO analysis (fold-change > 4; p < 0.05) suggested association of branch 2 with cell fate commitment towards the neuronal lineage, while branch 1 was associated with alternative developmental fates (Figures 5B-C; Table S7). Notably, no exogenous transcription factor was specifically enriched in branch 1 (Fisher’s exact test; p > 0.05) (Figures 5D and S5I). In contrast, branch 2 showed significant enrichment of 10 exogenous transcription factors (*ASCL1*, *DLX1, DLX2, FEV, FOXP2, ISL1, NEUROG2, NR4A2, PAX6* and *ZIC1;* p < 10^−2^). We independently transduced fibroblasts with the complete TF-pool (20 transcription factors), the 10 transcription factors enriched in branch 1, and the 10 transcription factors showing no enrichment in either branch. Neuronal profiling using immunofluorescence for TUBB3 and MAP2 and qPCR for pan-neuronal, neurotransmitter and fibroblast marker genes revealed that infection with transcription factors enriched in branch 2 markedly increased the efficiency of neuronal reprogramming compared to both the complete TF-pool and transcription factors that showed no branch-specific enrichment (Figures 5E-H). In fact, infection with transcription factors without branch-specific enrichment reduced the efficiency of neuronal conversion compared to the complete TF-pool and upregulated genes that were enriched in branch 1, such as *AREG* and *PTHR1*, during pseudo-temporal ordering (Figure 5I). Collectively, these data revealed that the cocktail of small molecules converted cells into “uncommited” cells, whereas combination of small molecules with ectopic expression of defined transcription factors enhanced the conversion towards the neuronal lineage.

**Figure 5.**
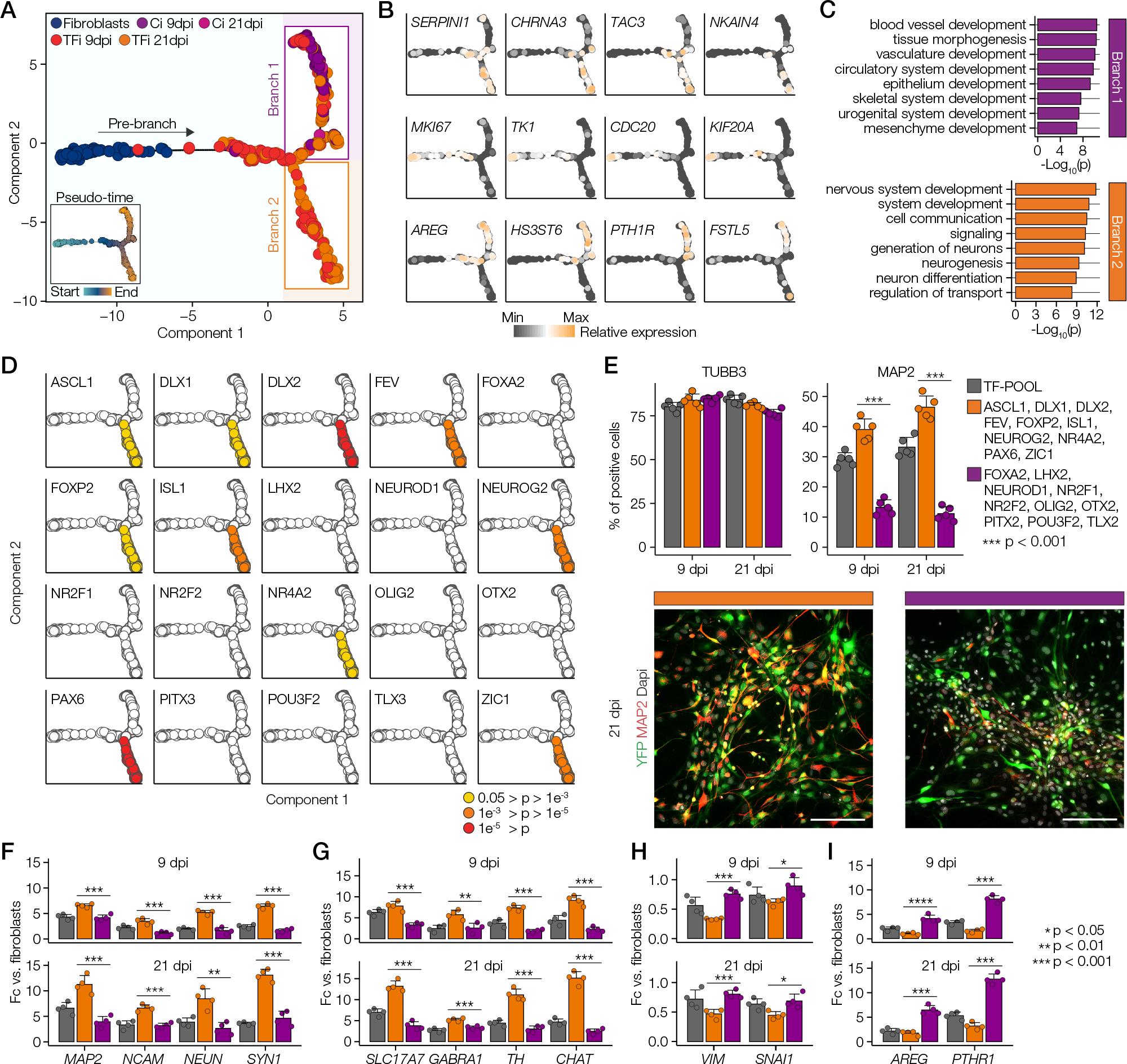
Association of developmental trajectories with exogenous expression profiles. (A) Pseudo-temporal ordering of cells (Fluidigm C1) based on the expression of 3,322 developmental genes (*n* = 446 cells). Small inset shows the same plot colored by pseudo-temporal values. (B) Branch-specific relative expression of neurogenic genes (top row), cell cycle-related genes (middle row) and genes related to alternative developmental fates (down row) along pseudo-time. (C) GO analysis of genes differentially expressed in branch 1 (top panel) and branch 2 (bottom panel). (D) Identification of exogenous transcription factors with branch-specific enrichment based on Fisher’s exact tests. (E) Top panels: Quantification of TUBB3+ and MAP2+ cells in fibroblasts infected with the complete TF-pool (gray), branch 2-enriched transcription factors (orange) and unenriched transcription factors (purple) at 9 dpi and 21 dpi. *n* = 5 independent experiments, error bars represent mean + SD. Bottom panels: Representative images of immunostainings for MAP2 (red) at 21 dpi; colors as in top panels. YFP (green) marks infected cells and cell nuclei were visualized using DAPI nuclear stain (grey). Scale bars, 100 μm. (F-I) Neuronal differentiation, loss of fibroblast characteristics and acquisition of alternative developmental fates as revealed by qPCR for pan-neuronal marker genes (*MAP2*, *NRCAM*, *NEUN*, *SYN1*; F), canonical neuronal subtype markers (*VGLUT1*, *GABA*, *TH*, *CHAT*; G), canonical fibroblast markers (*VIM*, *SNAI2*; H) and branch 1-enriched genes (*AREG*, *PTHR1*; I at 9 dpi and 21 dpi; colors as in E. *n* = 4 independent experiments, error bars represent mean + SD.

### Identification of GRNs controlling subtype specification of induced neurons

In previous reports, a single transcription factor was sufficient to reprogram cells to iN (Chanda et al., 2014; Liu et al., 2013). The reprogramming of the vast majority of neuronal subtypes, however, requires combinatorial expression of transcription factors in the same cell (Tsunemoto et al., 2018). To infer combinations of exogenous transcription factors governing neuronal subtype specification from our single-cell data, we classified each subtype based on canonical marker genes for glutamatergic, cholinergic, dopaminergic, GABAergic, serotonergic, glycinergic neurons, and neuronal progenitor cells (NPC). Next, exogenous transcription factors were assigned to each neuronal subtype network based on significant association with at least 3 endogenous marker genes (Fisher’s exact tests; p < 0.05) (Figure 6A). For NPC and glycinergic subtypes, only a single exogenous transcription factor (*ISL1* and *PAX6*, respectively) showed 3 or more associations, while no exogenous transcription factor showed more than 2 associations with serotonergic genes (Figure S6A). Notably, however, we identified various combinatorial enrichements of exogenous transcription factors associated with three or more of glutamatergic-, cholinergic-, dopaminergic- and GABAergic-related genes. Previous studies have shown that combination of *DLX1*, *DLX2* and *CTIP2* with miR-9/9*-124 converted human fibroblasts into an enriched population of neurons analogous to GABAergic striatal medium spiny neurons, whereas combination of *FOXA2*, *NR4A2* and *PITX3* with additional transcription factors generated dopaminergic neurons (Caiazzo et al., 2011; Liu et al., 2012; Pfisterer et al., 2011; Victor et al., 2014). Our results, in part, are consistent with these studies; two of the four transcription factors associating with the GABAergic GRN were *DLX1* and *DLX2*, and three of the five exogenous transcription factors associating with the dopaminergic GRN were *FOXA2*, *NR4A2* and *PITX3* (Figure 6B). Moreover, *ISL1*, which was previously combined with other transcription factors to convert human fibroblasts into cholinergic motor neurons (Son et al., 2011), associated stronger to the cholinergic GRN than to other GRNs. However, the distribution of transcription factors among subtype-specific GRNs was not mutually exclusive. For example, the glutamatergic GRN comprised transcription factors of all other GRNs, but further included *ASCL1*, *FEV*, *NEUROG2*. Thus, our data support a hierarchical reprogramming model which predicts that replacement of only a few factors can alter the fate of generated cells (Wapinski et al., 2013), and suggests that few transcription factors can override other transcription factors when co-expressed in the same cell.

**Figure 6.**
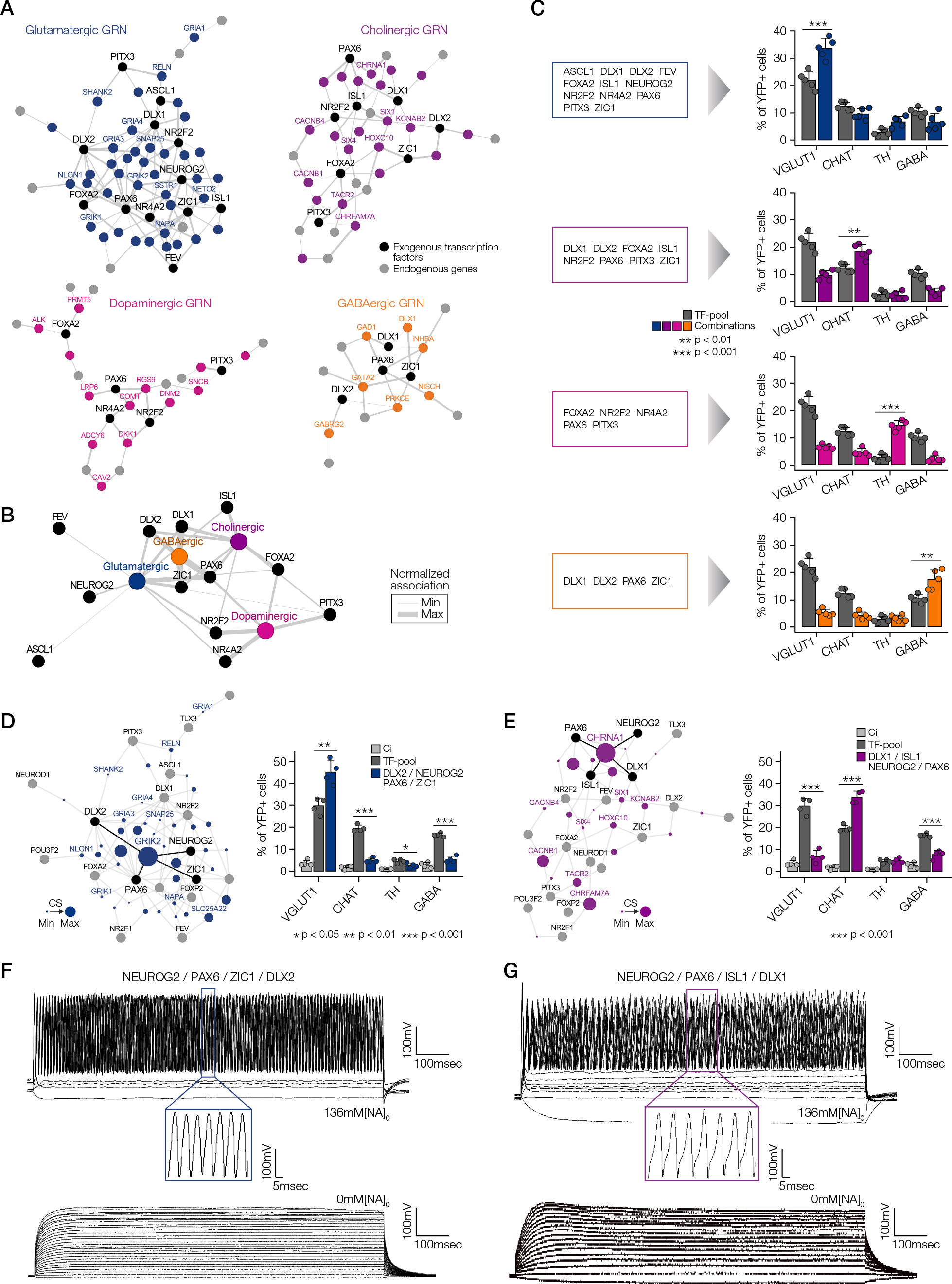
Identification of GRNs controlling subtype specification of induced neurons. (A) Neuronal subtype-specific gene regulatory networks on the basis of significant associations (Fisher’s exact test, p < 0.05). Neuronal subtype-specific genes are colored. Exogenous transcription factors with at least 3 edges to neuronal subtype-specific genes are shown in black. Exogenous transcription factors with less than 3 edges are shown in gray. (B) Edge-normalized network summarizing the associations of exogenous transcription factors (shown in A) with neuronal subtype-specific GRNs. (C) Validation of GRNs as revealed by quantifications of immunostainings for VGLUT1, CHAT, TH and GABA of fibroblasts separately infected with the TF-pool (gray) or with exogenous transcription factors shown in the boxes at 21 dpi (colored). *n* = 5 independent experiments, error bars represent mean + SD. (D-E) Left panels: visualization of Combinations Scores (CS) of glutamatergic-specific (D) and cholinergic-specific (E) GRNs. Exogenous transcription factors associated with genes showing highest CS in each gene regulatory network are shown in black, all other exogenous transcription factors are shown in gray. Neuronal subtype-specific genes are colored. Right panels: Validation of novel combinations of exogenous transcription factors by quantification of immunostainings for VGLUT1, CHAT, TH and GABA of Ci (light gray), fibroblasts infected with the complete TF-pool (dark gray) and fibroblasts infected with novel combinations (color) at 21 dpi. *n* = 4 independent experiments, error bars represent mean + SD. (F-G) The generation of the action potential in iN infected with *DLX2*, *NEUROG2*, *PAX6*, *ZIC1* (F) or *DLX1*, *ISL1*, *NEUROG2*, *PAX6* (G). Representative traces in the presence (upper panel) or absence (lower panel) of extracellular Na^+^ were recorded using the current-clamp protocol.

Subsequently, we independently tested all four predicted neuronal subtype combinations and evaluated expression of VGLUT1, CHAT, TH and GABA in reprogrammed neurons (Figures 6C and S6B). In all combinations analyzed, ectopic expression of predicted combinations significantly enriched for the intended target subset when compared to the complete TF-pool control (p < 0.01).

To identify minimal and robust combinations of transcription factors in glutamatergic and cholinergic GRNs, we attributed all genes Combination Scores (CS; Supplementary Methods). The CS represents the significance of cumulative marker gene expression in cells containing at least 2 of the predicted exogenous transcription factors (Figures 6D-E). In a second line of validation, we chose the gene with the highest CS for each network (*GRIK2* for the glutamatergic network and *CHRNA1* for the cholinergic network) and infected fibroblasts with predicted combinations of exogenous transcription factors. We evaluated phenotypes of iN at 21 dpi using immunofluorescence for VGLUT1, CHAT, TH and GABA. Infection with *GRIK2*-associated (*DLX2*, *NEUROG2*, PAX6, ZIC1) and *CHRNA1*-associated (*DLX1*, *ISL1*, *NEUROG2*, *PAX6*) transcription factors significantly enriched for glutamatergic and cholinergic neurons, respectively, when compared to fibroblasts infected with the complete TF-pool and Ci. Notably, PAX6, which associated with all four neuronal subtype-specific GRNs (Figures 6A-B), also was included in each of the newly identified combinations, indicating that similar to its role during normal neuronal development (Stoykova et al., 2000; Yun et al., 2001), PAX6 acts as a master regulator during neuronal reprogramming.

Finally, to validate the functionality of our induced glutamatergic and cholinergic neurons, we characterized the electrophysiological properties of iN infected with *DLX2*, *NEUROG2*, *PAX6*, *ZIC1* (Figure 6F) and *DLX1*, *ISL1*, *NEUROG2*, *PAX6* (Figure 6G). We observed that iN infected with both combinations of transcription factors could generate repetitive action potentials upon current injection (Figure 6F-G). We confirmed that these action potentials were inhibited by withdrawal of extracellular Na^+^. Notably, we observed that glutamatergic and cholinergic iN fired action potentials exhibiting different wave forms and showed significant differences in resting membrane potential and action potential duration, suggesting that the infection of the two different combinations of transcription factors generated iN with distinct electrophysiological properties.

Collectively, these results demonstrate that Convert-seq enabled efficient identification of GRNs and defined transcription factors directing the reprogramming of fibroblasts towards distinct, functional neuronal subtypes.

## Discussion

We developed Convert-seq, a method based on a previously published nested-PCR-based single-cell screening system (Shin et al., 2012), which adapts retrospective identification of vector-based gene expression in single-cells for the scale of massively-parallel single-cell RNA sequencing. Convert-seq overcomes the laborious and time-consuming process of identifying specific combinations of transcription factors capable to reprogram the original transcriptional network(Vierbuchen and Wernig, 2011). Particularly in less well-characterized cell types, it is a daunting task to test all possible combinations of candidate reprogramming factors in a one-by-one approach. The intricacy to disentangle the roles of exogenous and endogenous genes during reprogramming further complicates matters, which is why, despite considerable success in the reprogramming of various cell types, still little is known about gene regulatory networks (GRNs) controlling the reprogramming process(Gascon et al., 2017).

We have demonstrated that when combined with SMART-seq technology (Picelli et al., 2014; Ramskold et al., 2012), Convert-seq can reliably discriminate between endogenous and exogenous transcripts both at the bulk and single-cell level. Thus, this method allows to analyze transcriptional effects initiated by different combinations of candidate reprogramming factors in a single experiment. Since the number of single cells required to reach statistical significance grows exponentially as the number of candidate genes increases, Convert-seq is incapable of profiling all possible combinations. Therefore, we have limited our study to a subset of possible combinations by adjusting the multiplicity of infection (MOI) and culture conditions that favour the survival of neuronal cells (i.e. positive selection). Implementing Convert-seq in higher throughput platforms such as droplet-based scRNA-seq will further strengthen the utility of this approach. We tested Convert-seq in a droplet based 3‘-seq platform, however, detection of exogenous transcripts was inefficient, possibly due to the strong 3‘ bias of the protocol. Future work could provide droplet-based scRNA-seq platforms with more even coverage across transcripts or encode the identity of exogenous genes in sequence barcodes at the 3‘-end of constructs (Adamson et al., 2016; Dixit et al., 2016; Jaitin et al., 2016).

Achieving efficient generation of specific cell types still constitutes a central challenge in the field of reprogramming. However, using Convert-seq, we identified distinct developmental trajectories emerging early during the reprogramming of human fibroblasts to induced neurons and identified a set of 10 exogenous transcription factors to repress alternative developmental programs. Moreover, by discriminating exogenous and endogenous genes in single-cells, Convert-seq identified distinct combinations of transcription factors controlling the reprogramming of fibroblasts into different neuronal subtypes exhibiting distinct electrophysiological properties.

In addition to providing alternative sources for cell-based therapies, cell reprogramming could help to decipher the molecular logic behind subtype specification during normal development. Strikingly, our data recapitulates some of the general principles of normal neuronal development. For example, PAX6, a paired box transcription factor that acts as master regulator of the mammalian nervous system development and is expressed in region-specific manner in neuronal progenitor cells (NPCs) and uniformely in neuroectodermal cells differentiated from embryonic stem cells (ESCs) and iPSCs (Chapouton et al., 1999; Stoykova et al., 2000; Yun et al., 2001; Zhang et al., 2010), associated with all neuronal subtypes. *PAX6*-expressing neuroectodermal cells can be readily patterned to region specific NPCs that give rise to various neuronal subtypes including cholinergic neurons (Li et al., 2005), dopaminergic neurons (Yan et al., 2005) and GABAergic neurons (Kallur et al., 2008). Thus, our findings indicate that *PAX6* acts as a master regulator during reprogramming of iNs that, when combined with additional transcription factors, can instruct the specification into diverse neuronal subtypes. In line with this, our results show that combination of *PAX6* with *NEUROG2*, *DLX2* and *ZIC1* generates mainly glutamatergic neurons, whereas substitution of *DLX2* and *ZIC1* with *DLX1* and *ISL1* generates mainly cholinergic neurons. These findings are partly consistent with previous studies; in the dorsal telencephalon, *PAX6* and *NEUROG2* are involved in the specification of glutamatergic projection neurons, and overexpression of *Neurog2* in mouse embryonic stem cells and cultured mouse cortical astroglia instructs the reprogramming of functional glutamatergic neurons (Berninger et al., 2007; Heinrich et al., 2010; Schuurmans and Guillemot, 2002; Thoma et al., 2012). Conversely, combination of *NEUROG2* expression with specific small molecules converts human fibroblasts to cholinergic neurons (Liu et al., 2013; Smith et al., 2016). Additional studies will be required to determine if *PAX6*, analogous to the master regulator *ASCL1*, acts as pioneer factor that can access its targets even if they are bound by nucleosomes (Wapinski et al., 2013).

In conclusion, the data obtained by Convert-seq corroborates previous studies on neuronal reprogramming and predicts novel combinations of pro-neuronal transcription factors that allow the generation of clinically relevant neuronal subtypes. Thus, Convert-seq represents an effective method to identify transcriptional codes controlling multiple cell fate conversions, speaking to its potential to become a standard strategy to unravel molecular mechanisms governing cell reprogramming.

## Acknowledgments

This work was supported by a Research Grant from the Japanese Ministry of Education, Culture, Sports, Science and Technology (MEXT) and a Postdoctoral Fellowship for Research in Japan by the Japan Society for the Promotion of Science (JSPS). The authors wish to acknowledge RIKEN GeNAS for the sequencing of the libraries.

## Author Contributions

Formulation of research goals and aims: JL, JWS

Data curation: JL, TK, CP

Formal analysis: JL, TK, RN, JWS

Funding acquisition: JL, PC, JWS

Performing experiments: JL, TK, RN, TC, DS, MK, FR

Project supervision: PC, JWS

Writing – original draft, review & editing: JL, JWS

## Declaration of Interests

All authors do not claim conflict of interest for this study.

## STAR Methods

### Key Resources Table

| REAGENT or RESOURCE | SOURCE | IDENTIFIER |
| --- | --- | --- |
| **Antibodies** |  |  |
| Mouse monoclonal anti-Tubulin, beta, Class III (Tuj1) | Covance | Cat# MMS-435P;<br>RRID:AB_2313773 |
| Mouse monoclonal anti-Map2 | Abcam | Cat# ab11267;<br>RRID:AB_297885 |
| Rabbit polyclonal anti-Synapsin1 | Abcam | Cat# ab64581;<br>RRID:AB_1281135 |
| Rabbit polyclonal anti-vGlut1 | Synaptic Systems | Cat# 135 303<br>RRID:AB_887875 |
| Mouse monoclonal anti-GABA | Abcam | Cat# ab86186<br>RRID:AB_1925012 |
| Rabbit polyclonal anti-TH | Abcam | Cat# ab112<br>RRID:AB_297840 |
| Sheep polyclonal anti-CHAT | Abcam | Cat# ab18736<br>RRID:AB_2244867 |
| Goat anti-mouse IgG1 | Molecular Probes | Cat# A-21123<br>RRID:AB_141592 |
| Goat anti-mouse IgG2a | Thermo Fisher Scientific | Cat# A-21133<br>RRID:AB_2535772 |
| Goat polyclonal anti-rabbit IgG (H+L) | Thermo Fisher Scientific | Cat# R37117<br>RRID:AB_2556545 |
| Donkey polyclonal anti-sheep IgG (H+L) | Thermo Fisher Scientific | Cat# A-21098<br>RRID:AB_2535752 |
| **Chemicals, Peptides and Recombinant Proteins** |  |  |
| Fetal Bovine Serum (FBS) | Clontech | Cat# 631367 |
| Penicillin&Streptomycin | Thermo Fisher Scientific | Cat# 15140122 |
| Trypsin-EDTA | Sigma-Aldrich | Cat# 25200-114 |
| Polybrene | Sigma-Aldrich | Cat# H9268 |
| Poly-D-Lysine | Sigma-Aldrich | Cat# P6407 |
| Laminin from Engelbreth-Holm-Swarm murine sarcoma basement membrane | Sigma-Aldrich | Cat# L2020 |
| B27 Supplement (50x), serum free | Thermo Fisher Scientific | Cat# 17504044 |
| N2 Supplement (100x) | Thermo Fisher Scientific | Cat# 17502048 |
| MEM Non-Essential Amino Acids Solution (100x) | Thermo Fisher Scientific | Cat# 11140050 |
| Glutamax Supplement | Thermo Fisher Scientific | Cat# 35050061 |
| Valproic Acid for Biochemistry | WAKO | Cat# 227-01071 |
| L-Ascorbic Acid | Sigma-Aldrich | Cat# A92902 |
| ROCK inhibitor Y-27632 | WAKO | Cat# 253-00513 |
| N<sup>6</sup>,2'-O-Dibutyryladenosine 3',5'-cyclic monophosphate sodium salt | Sigma-Aldrich | Cat# D0260 |
| CHIR99021 | Abcam | Cat# ab120890 |
| SB-431542 | Sigma-Aldrich | Cat# S4317 |
| LDN-193189 | REPROCELL | Cat# 04-0074-10 |
| Human Noggin | Sigma-Aldrich | Cat# SRP4675 |
| BDNF Recombinant human protein | Thermo Fisher Scientific | Cat# 10908010 |
| NT3 Recombinant human protein | R&D Systems | Cat# 267-N3 |
| GNDF Recombinant human protein | Thermo Fisher Scientific | Cat# PHC7044 |
| Proteinase K Solution | Thermo Fisher Scientific | Cat# AM2548 |
| FuGENE HD Transfection Reagent | Promega | Cat# E2311 |
| HBSS, calcium, magnesium, no phenol red | Thermo Fisher Scientific | Cat# 14025092 |
| RNase ONE Ribonuclease | Promega | Cat# M4261 |
| Triton X-100 | Sigma-Aldrich | Cat# T8787 |
| **Critical Commercial Assays** |  |  |
| Gateway LR Clonase II Enzyme mix | Thermo Fisher Scientific | Cat# 11791100 |
| PureYield Plasmid Midiprep System | Promega | Cat# A2492 |
| RNeasy Mini Kit | Quiagen | Cat# 74106 |
| One-Step SYBR PrimeScript RT PCR Kit II | Takara | Cat# RR086A |
| Chromium™ Single Cell 3' Reagent kits V1 | 10x Genomics | Cat# CG00026 |
| KAPA Library Quantification kit | KAPA BIOSYSTEMS | Cat# 07960166001 |
| ERCC RNA Spike-In Mix1 | Thermo Fisher Scientific | Cat# 4456740 |
| SMARTer Ultra Low RNA Kit | Clontech | Cat# 634848 |
| Quant-iT™ PicoGreen dsDNA Assay kit | Thermo Fisher Scientific | Cat# P7589 |
| Nextera XT DNA Library Preparation kit | Illumina | Cat# FC-131-1024 |
| Nextera XT Index Kit v2 | Illumina | Cat# FC-131-1002 |
| **Deposited Data** |  |  |
| Raw and analyzed data | This Paper | GEO: GSE117075 |
| **Experimental Models: Cell Lines** |  |  |
| Human neonatal dermal fibroblasts | Lonza | Cat# C-2509 |
| HEK293T | ATCC | Cat# CRL-3216 |
| **Recombinant DNA** |  |  |
| CSII-EF-RfA-IRES2-VENUS-ASCL1 | This paper | N/A |
| CSII-EF-RfA-IRES2-VENUS-DLX1 | This paper | N/A |
| CSII-EF-RfA-IRES2-VENUS-DLX2 | This paper | N/A |
| CSII-EF-RfA-IRES2-VENUS-FEV | This paper | N/A |
| CSII-EF-RfA-IRES2-VENUS-FOXA2 | This paper | N/A |
| CSII-EF-RfA-IRES2-VENUS-FOXP2 | This paper | N/A |
| CSII-EF-RfA-IRES2-VENUS-ISL1 | This paper | N/A |
| CSII-EF-RfA-IRES2-VENUS-LHX2 | This paper | N/A |
| CSII-EF-RfA-IRES2-VENUS-NEUROD1 | This paper | N/A |
| CSII-EF-RfA-IRES2-VENUS-NEUROG2 | This paper | N/A |
| CSII-EF-RfA-IRES2-VENUS-NR2F1 | This paper | N/A |
| CSII-EF-RfA-IRES2-VENUS-NR2F2 | This paper | N/A |
| CSII-EF-RfA-IRES2-VENUS-NR4A2 | This paper | N/A |
| CSII-EF-RfA-IRES2-VENUS-OLIG2 | This paper | N/A |
| CSII-EF-RfA-IRES2-VENUS-OTX2 | This paper | N/A |
| CSII-EF-RfA-IRES2-VENUS-PAX6 | This paper | N/A |
| CSII-EF-RfA-IRES2-VENUS-PITX3 | This paper | N/A |
| CSII-EF-RfA-IRES2-VENUS-POU3F2 | This paper | N/A |
| CSII-EF-RfA-IRES2-VENUS-TLX3 | This paper | N/A |
| CSII-EF-RfA-IRES2-VENUS-ZIC1 | This paper | N/A |
| **Software and Algorithms** |  |  |
| Bowtie v1.2.2 | (Langmead et al., 2009) | http://bowtie-bio.sourceforge.net/index.shtml |
| Kallisto v0.42.4 | (Bray et al., 2016) | https://pachterlab.github.io/kallisto/ |
| R v3.5.2 | R project | https://www.r-project.org/ |
| edgeR v3.24.3 | (Robinson et al., 2010) | https://bioconductor.org/packages/release/bioc/html/edgeR.html |
| rio v0.5.16 | CRAN | https://cran.r-project.org/web/packages/rio/index.html |
| Tidyverse v1.2.1 | CRAN | https://cran.r-project.org/web/packages/tidyverse/index.html |
| Readxl v1.2.0 | CRAN | https://cran.r-project.org/web/packages/readxl/index.html |
| Reshape v0.8.8 | (Wickham, 2007) | https://cran.r-project.org/web/packages/reshape/index.html |
| pheatmap v1.0.12 | CRAN | https://cran.r-project.org/web/packages/pheatmap/index.html |
| data.table v1.12.0 | CRAN | https://cran.r-project.org/web/packages/data.table/index.html |
| Seurat v2.3.0 | (Butler et al., 2018) | https://cran.r-project.org/web/packages/Seurat/index.html |
| Dendextend v1.9.0 | (Galili, 2015) | https://cran.r-project.org/web/packages/dendextend/citation.html |
| monocle v2.2.0 | (Trapnell et al., 2014) | https://bioconductor.org/packages/release/bioc/html/monocle.html |
| Cytoscape v3.7.1 | (Shannon et al., 2003) | https://cytoscape.org/ |

### Cell culture and generation of iN

Human neonatal dermal fibroblasts were purchased from Lonza (C-2509; passages 4-8) and cultured in DMEM containing 10% FBS/L-glutamine/10% Penicilin&Streptomycin at 37°C 5% CO_2_ until virus infection. For TFi generation, a total of 6 MOI (~0.3 MOI per virus) were pooled into culture medium containing 8 µg*ml^−1^ Polybrene (SIGMA) to increase infection efficiency. The virus pool was infected and incubated overnight at 37°C 5% CO_2_. On the following day, virus-containing medium was removed and cells were incubated for an additional day in fresh culture medium at 37°C 5% CO_2_. For iN generation, infected cells were split onto Poly-d-lysine (100μg*ml^−1^; SIGMA) / Laminin (50μg*ml^−1^; SIGMA)-coated culture plates and incubated overnight in culture medium at 37°C 5% CO_2_. On the following day, medium was changed to neuronal induction medium containing DMEM/F12 and Neurobasal-A (Thermo Fisher Scientific) mixed at a 1:1 ratio, 2% (vol/vol) B27 supplement and 0.5% N2 Supplement (GIBCO), 1× nonessential amino acids (Thermo Fisher Scientific), 1% GlutaMAX supplement, VPA (0.5mM, WAKO), L-Ascorbic acid (200nM, SIGMA), Y-27632 (10μM, WAKO), dcAMP (0.5mM, SIGMA), 10% FBS and 10% Penicilin&Streptomycin (WAKO). The concentrations of small molecules used were as follows: CHIR99021 (2μM, Abcam), SB-431542 (10μM, SIGMA), LDN-193189 (0.5μM, Stemgent), Noggin (100ng*ml^−1^, SIGMA). Neuronal induction medium was changed every third day, during which the concentration of FBS was gradually reduced from 10% to 0%. After 2 weeks, neuronal induction medium was replaced with neuronal maturation medium without small molecules, but containing 10ng*ml^−1^ BDNF (GIBCO), 10ng*ml^−1^ NT3 (R&D Systems) and 10ng*ml^−1^ GDNF (Thermo Fisher Scientific) and the medium was changed every third day until further analysis.

### Complementary DNA and virus generation

Complementary DNA (cDNA) and viruses were generated as described previously (Shin et al., 2012). Briefly, we recombined Gateway-compatible human full-length cDNA entry clones derived from RIKEN BRC clone bank (http://www.brc.riken.jp/) into the pENTR lentivirus vector CSII-EF-RfA-IRES2-VENUS using Gateway LR clonase II enzyme mix (Invitrogen). After Proteinase K treatment, recombined plasmids were transformed into competent *Escherichia coli* and plasmids derived from single colonies were expanded and purified using PureYield Plasmid Midiprep System (Promega). Plasmids, HIV-gp and VSV envelope genes were co-transfected onto 293T cells using FuGeneHD (Roche). Supernatant-containing viruses were collected, centrifuged by ultracentrifugation and dissolved in 100µl HBSS buffer (WAKO) and stored at −80°C for later use.

### Immunocytochemistry and quantitative RT-PCR

For immunocytochemistry, cells were fixed in 4% paraformaldehyde for 20 min at room temperature and permeabilized using 0.2% Triton X-100 (SIGMA) for 10 min at room temperature. Following permeabilization, cells were pre-incubated with blocking solution (2% BSA, 0.2% Triton X-100) to block non-specific sites for 1 h. Primary antibodies were diluted in blocking solution and applied to cells over night at 4°C. Secondary antibodies were diluted in blocking solution and applied to cells at room temperature for 1 h. Imaging was performed using the INCell Analyzer 6000 (GE Healthcare). The following primary antibodies and dilutions were used: mouse anti-TUBB3 (Covance, 1:1000), mouse anti-MAP2 (Abcam, 1:500), rabbit anti-SYNAPSIN 1 (Abcam, 1:200), rabbit anti-VGLUT1 (Synaptic Systems, 1:100), mouse anti-GABA (Abcam, 1:200), sheep anti-CHAT (Abcam, 1:100), rabbit anti-TH (Abcam, 1:500). The following secondary antibodies and dilutions were used: goat anti-mouse IgG1 (GIBCO, 1:200), goat anti-mouse IgG2a (Termo Fisher Scientific, 1:200), goat anti-rabbid IgG (Thermo Fisher Scientific, 1:200), donkey anti-sheep IgG (Thermo Fisher Scientific, 1:200). Human neonatal dermal fibroblasts were used as negative controls. Quantification of immunostainings was performed using the INCell Investigator Developer Toolbox. For quantitative RT-PCR (qRT-PCR), total RNA was purified using the RNeasy Mini Kit (QIAGEN) according to the manufacturer’s specification. Quality and quantity of RNA was determined using a DropSense96 (Trinean). Equal amounts of RNA were reverse-transcribed using the One-Step SYBR PrimeScript RT PCR Kit II, and cDNAs were normalized to equal amounts using primers against *Gapdh*. qRT-PCR was performed on a 7900HT Fast Real-Time PCR system (Applied Biosystems).

### Convert-seq

#### Artificial transcript model

To determine the exact nucleotide sequences flanking the ORF of each exogenous transcript, we sequenced recombined plasmids using a 3730/3730xl DNA Analyzer (Applied Biosystems) following the manufacturer’s protocol. Briefly, we first amplified templates by PCR using a primers annealing to the EF1A promoter sequence near the 5‘ end of each ORF to amplify the 5‘ junction sequences and primers annealing to the IRES2 sequence near the 3‘ end of each ORF to amplify the 3‘ junction sequences. After gel purification, we sequenced templates using BigDye Terminator v3.1 Cycle Sequencing Kits (Applied Biosystems). We integrated results derived from 3 primers (3 replicates each) at both the 5‘ and 3‘ junction of each ORF and combined the resulting 5‘ and 3‘ junction sequences with known sequences of the ORFs of each transcription factor and the CSII-EF-RfA-IRES2-VENUS pENTR lentivirus vector to compile the artificial exogenous transcript model. Lastly, we combined our artificial exogenous transcript model with the human transcriptome (version GRCh38.p5) to obtain the final artificial transcript model.

#### Read alignment with Bowtie

Reads were aligned to the artificial transcript model using Bowtie v1.2.2 with the default parameter settings for paired-end reads. After retrieving BED12 files using samtools and bedtools, we intersected all reads using a custom GFF file in which 5‘ and 3‘ junctions of all exogenous sequences were defined. Only reads overlapping the junction sequences by at least 5 bp were counted as specific reads. The expression values of all exogenous transcription factors were quantified as count per million (CPM) and transformed to log^2^ (CPM+1).

#### Read alignment with Kallisto

For alignment using Kallisto (v0.42.4), alignment to the full artificial transcript model yielded many false-positive hits (Figure S3B). Therefore, we trimmed the 5‘ and 3‘ junction sequences to ~100 bp on either side, which markedly reduced the number of false positive hits (Figure 3D). Reads were aligned with the default parameter settings for paired-end reads. Custom R scripts were used to merge transcript isoforms and compile a single expression matrix.

#### Convert-seq on Drop-seq data

Since the CellRanger Software does not provide a single FASTQ file for each single-cell, we used the bcl2fastq Conversion Software to convert and demultiplex BCL files. After retrieving FASTQ files for each single cell, reads were aligned to the trimmed artificial transcript model using Kallisto with the default parameter settings in paired-end mode.

### Droplet-based scRNA-seq

#### Library preparation and sequencing

Droplet-based scRNA-seq libraries were generated using the Chromium^TM^ Single Cell 3’ Reagent kits V1 (CG00026, 10× Genomics). Briefly, cell number and cell viability were assessed using the Countess II Automated Cell Counter (ThermoFisher). Thereafter, cells were mixed with the Single Cell Master Mix and loaded together with Single Cell 3’ Gel beads and Partitioning Oil into a Single Cell 3’ Chip. RNA transcripts were uniquely barcoded and reverse-transcribed in droplets. cDNAs were pooled and amplified according to the manufacturer’s protocol. Libraries were quantified by High Sensitivity DNA Reagents (Agilent Technologies) and the KAPA Library Quantification kit (KAPA BIOSYSTEMS). Libraries then were sequenced by Illumina Hiseq 2500 in rapid mode.

#### Read alignment and gene quantification

Initial read alignment to hg19 human reference genome, filtering and UMI counting was performed by the CellRanger Software ver 1.1.0 using default parameters. This software implements STAR as an alignment tool. Data from TFi and Ci were normalized to the same sequencing depth and aggregated into a single gene-barcode matrix. The expression values were quantified as count per million (CPM) and transformed to log^2^ (CPM+1).

### scRNAseq using the Fluidigm C1 platform

#### Library preparation and sequencing

Single cell RNA-seq analysis was performed following the manufacturer’s protocol (P100-7168L1, Fluidigm). Briefly, cell number and cell viability were assessed using the Countess II Automated Cell Counter (ThermoFisher). After priming medium size C1 Single-cell Open App IFCs, 250 cells/μL were loaded and capture efficiency and cell morphology was assessed using the IN Cell Analyzer 6000 (GE Healthcare). To exclude chambers loaded with no cells, more than one cell (cell doublets) or dead cells for downstream analysis, we took 11 z-stacking images per chamber. Next, cells were lysed with 20,000-fold diluted ERCC RNA Spike-In Mix1 (Thermo Fisher Scientific) and reverse transcription (RT) and cDNA amplification were performed using the SMARTer Ultra Low RNA Kit for the Fluidigm C1^TM^ System (Clontech). The amplified cDNAs were harvested into 96 well plates and quantified with Quant-iT ^TM^ PicoGreen dsDNA Assay kit. Library preparation was performed with the Nextera XT DNA Library Preparation kit (Illumina), Nextera XT Index Kit v2 (Illumina) and AMpure XP beads (Beckman Coulter). Libraries were quantified by High Sensitivity DNA Reagents (Agilent Technologies) and KAPA Library Quantification kit (KAPA BIOSYSTEMS). Each of the libraries were sequenced by Illumina Hiseq 2500 in highoutput mode (100bp paired end).

#### Read alignment and gene quantification

Reads were aligned to the trimmed artificial transcript model using Kallisto with the default parameter settings for paired-end reads. The expression values were quantified as transcript per million (TPM) and transformed to log^2^ (TPM+1).

### Fluidigm C1 reversed loading protocol (backloading) for bulk RNAseq

To perform bulk RNAseq of a total of 96 samples, we used the Fluidigm Script Builder^TM^ to design a reversed protocol that allows to load each sample into a separate chamber, where RT and cDNA amplification is performed. After priming the chips, 25 ng of RNA of each sample was loaded into the output wells on a medium size C1 Single-cell Open App IFC and the IFC was sealed using a C1 Porous Barrier Tape kit (Fluidigm). RT and cDNA amplification was performed following the manufacturer’s protocol (P100-7168L1). We ran the backloading script for 15 min at 4°C and switched to the mRNA seq RT and Amp script (1772×), which harvested cDNA back into the output wells. To remove remaining RNA, we added Rnase One Ribonuclease (Promega) at room temperature. To quantify the cDNA, we used the Quant-iT PicoGreen dsDNA Assay kit. Library preparation was performed using the Nextera XT DNA Library Preparation kit (Illumina), the Nextera XT Index Kit v2 (Illumina) and Ampure XP beads (Beckman Coulter). Libraries were quantified using the High Sensitivity DNA Reagents (Agilent Technologies) and the KAPA Library Quantification kit (KAPA BIOSYSTEMS). Libraries were sequenced on the Illumina Hiseq 2500 platform in rapid mode (100bp paired end).

### Electrophysiology

Conventional whole-cell current-clamp recordings were performed as described previously (Ichikawa et al., 2012). All experiments were conducted at 25°C. Patch pipettes with resistances ranging from 3–7 MΩ were pulled from capillary tubes using a DMZ-Universal puller (Zeitz Instruments GmbH, Martinsried, Germany) and then back-filled with intracellular solution. Action potentials were recorded by a patch clamp amplifier (axopatch 200B; Axon Instruments, Foster City, CA) with a series of current step from 0 to 200 pA with a 2,000-ms duration. The action potentials were monitored and stored using pCLAMP software (Molecular Devices, LLC., CA) after digitizing the analog signals at 5 kHz (DigiData 1322A; Axon Instruments, Foster City, CA). For patch-clamp recordings, the extracellular solution (ECS) consisted of the following: 137 mM NaCl, 5 mM KCl, 0.44 mM KH_2_PO_4_, 0.33 mM Na_2_HPO_4_, 10 mM glucose, 12 mM NaHCO_3_, 0.5 mM MgCl_2_, and 10 mM HEPES, adjusted to pH 7.4 with tris(hydroxymethyl)aminomethane. To examine the Na^+^ selectivity, extracellular 136 mM NaCl was substituted with equimolar extracellular LiCl (Na^+^-free ECS). To record ionic currents under physiological conditions, intracellular solution containing 150 mM KCl, 10 mM HEPES, and 2 mM magnesium adenosine triphosphate (pH 7.2 by tris(hydroxymethyl)aminomethane) was used.

### Computational methods for scRNA-seq data

#### Quality control, cell clustering and t-SNE visualization

All analyzes and visualization of data were conducted in an R environment. Clustering and t-SNE visualization was performed using the R package ‘Seurat’ (Satija et al., 2015) (v2.3.0). For droplet-based scRNA-seq data, genes expressed in less than 3 cells and cells expressing less than 1000 genes or more than 4500 genes were removed. In addition, we removed cells expressing more than 2% mitochondrial genes, indicative of dead cells. PCA was performed on the ~1000 most variable genes after regressing out the number of UMI and the percentage of mitochondrial genes. Using the 20 most significant principal components (PCs), we projected individual cells based on their PC scores onto a single two-dimensional map using t-SNE (Van Der Maaten and Hinton, 2008). Gene expression heatmap along t-SNE2 was obtained by dividing cells into 40 groups based on their t-SNE2 scores, averaging gene expression within each group and scaling expression values by column. For Fluidigm C1 data, we excluded chambers containing no cells, multiple cells or cells exhibiting morphological features of cell death based on visual inspection using the IN Cell Analyzer 6000 (GE Healthcare). Additionally, cells expressing either of the two housekeeping genes Actb and Gapdh (encoding β-actin and glyceraldehyde-3-phosphate dehydrogenase, respectively) at less than three standard deviations below the mean were scored as unhealthy and removed. After applying these filters, 78 fibroblasts, 216 cells for the time-point 9 dpi (87 Ci and 129 TFi) and 152 cells for the time-point 21 dpi (15 Ci and 137 TFi) remained, yielding 446 cells in total. Genes expressed in less than 3 cells were removed. PCA was performed on the ~5000 most variable genes. Using the 5 most significant principal components (PCs), we projected individual cells based on their PC scores onto a single two-dimensional map using t-SNE. Hierarchical clustering was performed on cells and on PCA scores using Euclidean distance metric.

#### Differential expression test and GO analysis

Marker genes of each cluster were determined using a likelyhood ratio test based on zero-inflated data (p < 1e-4). We used marker genes which showed, on average, at least 3-fold and 2-fold enrichment (droplet-based scRNA-seq data and Fluidigm C1 data, respectively) in a cluster compared to all other clusters. GO analysis was performed using the PANTHER data base (http://www.pantherdb.org/) which uses Fisher’s Exact tests with FDR multiple test correction.

#### Construction of cellular network

The cellular network was constructed by computing a pairwise correlation matrix of all cells in our time-course data and the primary cortical and medial ganglionic eminence cells (Nowakowski et al., 2017). Next, we generated a weighted adjacency network graph using the perfuse force-directed layout in Cytoscape and visualized cells as nodes connected to other cells via edges if the Pearson pairwise correlation between two cells was >= 0.4.

#### Pseudotemporal ordering

Pseudotemporal ordering of cells was performed using the R package ‘Monocle’ (Trapnell et al., 2014) (v2.2.0). For unsupervised ordering, we used genes differentially expressed between cells at day 0 (fibroblasts) and Ci and TFi at day 9 and day 21 (qval < 0.1; ~10‘000 genes). To determine genes that are significantly branch-dependent (p < 10^−4^), we applied the BEAM algorithm. GO analysis for branch-dependent genes was performed using genes that met the following criteria: 1) p < 0.01 in a likelyhood ratio test based on zero-inflated data; 2) absolute log_2_ fold changes between the branch under consideration and others were larger than 2. GO analysis for genes that changed significantly as a function of pseudotime was performed using genes that met the following criteria: 1) p < 10^−4^ of differentialGeneTest; 2) among the top 1000 genes showing positive or negative correlation with pseudotime values. For semi-supervised ordering, we used ~3000 genes previously implicated in nervous system development (GO:0007399), circulatory system development (GO:0072359), urogenital system development (GO:0001655), heart development (GO:0007507), mesenchyme development (GO:0060485), ear development (GO:0043583), muscle structure development (GO:0061061), stem cell development (GO:0048864), pancreas development (GO:0031016) and skeletal system development (GO:0001501). GO analysis was performed using genes that met the following criteria: 1) p < 0.01 in a likelyhood ratio test based on zero-inflated data; 2) absolute log^2^ fold changes between the branch under consideration and others were larger than 2. To determine exogenous transcription factors that are significantly branch dependent, expression values were binarized (0 = not expressed, 1 = expressed). Then we performed Fisher’s exact tests to calculate the significance of association of a given exogenous TF with each cluster. Exogenous transcription factors with p < 0.05 were considered significantly enriched.

#### GRN assembly

To construct neuronal subtype-specific GRNs, we calculated the significance of association of each exogenous transcription factors with all other genes using Fisher‘s exact tests. Exogenous TFs were attributed to neuronal subtype-specific GRNs based on the following criteria: 1) p < 0.05 in Fisher‘s exact test; 2) at least 3 edges to 3 neuronal subtype-specific genes. GRNs were visualized with Cytoscape using the organic layout.

#### Combination score

The CS represents the -log10-transformed p-value of the significance (Mann-Whitney U test) of increased gene expression in cells containing at least 2 of the predicted exogenous transcription factors in a network compared to all other cells.

### Statistics

Statistical analyses were performed using R and detailed in the corresponding figure legends. All Stundent’s *t*-tests are two-sided.

### Code availability

All analysis code used in this study is available upon request. Custom code for the main analytical steps will be uploaded on GitHub at the day of publication.

### Data availability

All sequence data used in this study have been deposited in the NCBI Gene Expression Omnibus database and will be accessible through accession number GSE117075 upon publication.

## References

Adamson, B., Norman, T.M., Jost, M., Cho, M.Y., Nunez, J.K., Chen, Y., Villalta, J.E., Gilbert, L.A., Horlbeck, M.A., Hein, M.Y., et al. (2016). A Multiplexed Single-Cell CRISPR Screening Platform Enables Systematic Dissection of the Unfolded Protein Response. Cell 167, 1867–1882 e21.

Berninger, B., Costa, M.R., Koch, U., Schroeder, T., Sutor, B., Grothe, B., and Gotz, M. (2007). Functional Properties of Neurons Derived from In Vitro Reprogrammed Postnatal Astroglia. J. Neurosci. 27, 8654–8664.

Bray, N.L., Pimentel, H., Melsted, P., and Pachter, L. (2016). Near-optimal probabilistic RNA-seq quantification. Nat Biotechnol 34, 525–527.

Butler, A., Hoffman, P., Smibert, P., Papalexi, E., and Satija, R. (2018). A n a ly s i s Integrating single-cell transcriptomic data across different conditions, technologies, and species. 36.

Caiazzo, M., Dell’Anno, M.T., Dvoretskova, E., Lazarevic, D., Taverna, S., Leo, D., Sotnikova, T.D., Menegon, A., Roncaglia, P., Colciago, G., et al. (2011). Direct generation of functional dopaminergic neurons from mouse and human fibroblasts. Nature 476, 224–227.

Chanda, S., Ang, C.E., Davila, J., Pak, C., Mall, M., Lee, Q.Y., Ahlenius, H., Jung, S.W., Sudhof, T.C., and Wernig, M. (2014). Generation of induced neuronal cells by the single reprogramming factor ASCL1. Stem Cell Reports 3, 282–296.

Chapouton, P., Gartner, A., and Gotz, M. (1999). The role of Pax6 in restricting cell migration between developing cortex and basal ganglia. Development 126, 5569–5579.

Chen, S., Sanjana, N.E., Zheng, K., Shalem, O., Lee, K., Shi, X., Scott, D.A., Song, J., Pan, J.Q., Weissleder, R., et al. (2015). Genome-wide CRISPR screen in a mouse model of tumor growth and metastasis. Cell 160, 1246–1260.

Dixit, A., Parnas, O., Li, B., Chen, J., Fulco, C.P., Jerby-Arnon, L., Marjanovic, N.D., Dionne, D., Burks, T., Raychowdhury, R., et al. (2016). Perturb-Seq: Dissecting Molecular Circuits with Scalable Single-Cell RNA Profiling of Pooled Genetic Screens. Cell 167, 1853–1866 e17.

Galili, T. (2015). Data and text mining dendextend: an R package for visualizing, adjusting and comparing trees of hierarchical clustering. 31, 3718–3720.

Gascon, S., Masserdotti, G., Russo, G.L., and Gotz, M. (2017). Direct Neuronal Reprogramming: Achievements, Hurdles, and New Roads to Success. Cell Stem Cell 21, 18–34.

Heinrich, C., Blum, R., Gascón, S., Masserdotti, G., Tripathi, P., Sánchez, R., Tiedt, S., Schroeder, T., Götz, M., and Berninger, B. (2010). Directing astroglia from the cerebral cortex into subtype specific functional neurons. PLoS Biol. 8.

Hsu, P.D., Lander, E.S., and Zhang, F. (2014). Development and applications of CRISPR-Cas9 for genome engineering. Cell 157, 1262–1278.

Hu, W., Qiu, B., Guan, W., Wang, Q., Wang, M., Li, W., Gao, L., Shen, L., Huang, Y., Xie, G., et al. (2015). Direct Conversion of Normal and Alzheimer’s Disease Human Fibroblasts into Neuronal Cells by Small Molecules. Cell Stem Cell 17, 204–212.

Jaitin, D.A., Weiner, A., Yofe, I., Lara-Astiaso, D., Keren-Shaul, H., David, E., Salame, T.M., Tanay, A., van Oudenaarden, A., and Amit, I. (2016). Dissecting Immune Circuits by Linking CRISPR-Pooled Screens with Single-Cell RNA-Seq. Cell 167, 1883–1896 e15.

Jeff, G., Schnapp, B.J., and Sheetz, M.P. (1988). © 198 8 Nature Publishing Group. Nature 331, 450.

Jiang, H., Xu, Z., Zhong, P., Ren, Y., Liang, G., Schilling, H.A., Hu, Z., Zhang, Y., Wang, X., Chen, S., et al. (2015). Cell cycle and p53 gate the direct conversion of human fibroblasts to dopaminergic neurons. Nat. Commun. 6, 1–14.

Kallur, T., Gisler, R., Lindvall, O., and Kokaia, Z. (2008). Pax6 promotes neurogenesis in human neural stem cells. Mol Cell Neurosci 38, 616–628.

Kim, J., Su, S.C., Wang, H., Cheng, A.W., Cassady, J.P., Lodato, M.A., Lengner, C.J., Chung, C.Y., Dawlaty, M.M., Tsai, L.H., et al. (2011). Functional integration of dopaminergic neurons directly converted from mouse fibroblasts. Cell Stem Cell 9, 413–419.

Langmead, B., Trapnell, C., Pop, M., and Salzberg, S.L. (2009). Ultrafast and memory-efficient alignment of short DNA sequences to the human genome. Genome Biol 10, R25.

Li, X., Zuo, X., Jing, J., Ma, Y., Wang, J., Liu, D., Zhu, J., Du, X., Xiong, L., Du, Y., et al. (2015). Small-Molecule-Driven Direct Reprogramming of Mouse Fibroblasts into Functional Neurons. Cell Stem Cell 17, 195–203.

Li, X.J., Du, Z.W., Zarnowska, E.D., Pankratz, M., Hansen, L.O., Pearce, R.A., and Zhang, S.C. (2005). Specification of motoneurons from human embryonic stem cells. Nat Biotechnol 23, 215–221.

Liu, M.L., Zang, T., Zou, Y., Chang, J.C., Gibson, J.R., Huber, K.M., and Zhang, C.L. (2013). Small molecules enable neurogenin 2 to efficiently convert human fibroblasts into cholinergic neurons. Nat Commun 4, 2183.

Liu, M.L., Zang, T., and Zhang, C.L. (2016). Direct Lineage Reprogramming Reveals Disease-Specific Phenotypes of Motor Neurons from Human ALS Patients. Cell Rep. 14, 115–128.

Liu, X., Li, F., Stubblefield, E.A., Blanchard, B., Richards, T.L., Larson, G.A., He, Y., Huang, Q., Tan, A.C., Zhang, D., et al. (2012). Direct reprogramming of human fibroblasts into dopaminergic neuron-like cells. Cell Res. 22, 321–332.

Liu, Y., Yu, C., Daley, T.P., Wong, W.H., Wernig, M., and Qi, L.S. (2018). Resource CRISPR Activation Screens Systematically Identify Factors that Drive Neuronal Fate and Resource CRISPR Activation Screens Systematically Identify Factors that Drive Neuronal Fate and Reprogramming. Stem Cell 23, 758–771.e8.

Van Der Maaten, L.J.P., and Hinton, G.E. (2008). Visualizing high-dimensional data using t-sne. J. Mach. Learn. Res. 9, 2579–2605.

Masserdotti, G., Gascon, S., and Gotz, M. (2016). Direct neuronal reprogramming: learning from and for development. Development 143, 2494–2510.

Mazzoni, E.O., Mahony, S., Closser, M., Morrison, C.A., Nedelec, S., Williams, D.J., An, D., Gifford, D.K., and Wichterle, H. (2013). Synergistic binding of transcription factors to cell-specific enhancers programs motor neuron identity. Nat. Neurosci. 16, 1219–1227.

Mertens, J., Paquola, A.C.M., Ku, M., Hatch, E., Böhnke, L., Ladjevardi, S., McGrath, S., Campbell, B., Lee, H., Herdy, J.R., et al. (2015). Directly Reprogrammed Human Neurons Retain Aging-Associated Transcriptomic Signatures and Reveal Age-Related Nucleocytoplasmic Defects. Cell Stem Cell 17, 705–718.

Nowakowski, T.J., Bhaduri, A., Pollen, A.A., Alvarado, B., Mostajo-Radji, M.A., Di Lullo, E., Haeussler, M., Sandoval-Espinosa, C., Liu, S.J., Velmeshev, D., et al. (2017). Spatiotemporal gene expression trajectories reveal developmental hierarchies of the human cortex. Science (80-.). 358, 1318–1323.

Pang, Z.P., Yang, N., Vierbuchen, T., Ostermeier, A., Fuentes, D.R., Yang, T.Q., Citri, A., Sebastiano, V., Marro, S., Sudhof, T.C., et al. (2011). Induction of human neuronal cells by defined transcription factors. Nature 476, 220–223.

Pfisterer, U., Kirkeby, A., Torper, O., Wood, J., Nelander, J., Dufour, A., Bjorklund, A., Lindvall, O., Jakobsson, J., and Parmar, M. (2011). Direct conversion of human fibroblasts to dopaminergic neurons. Proc. Natl. Acad. Sci. 108, 10343–10348.

Picelli, S., Faridani, O.R., Bjorklund, A.K., Winberg, G., Sagasser, S., and Sandberg, R. (2014). Full-length RNA-seq from single cells using Smart-seq2. Nat Protoc 9, 171–181.

Rackham, O.J., Firas, J., Fang, H., Oates, M.E., Holmes, M.L., Knaupp, A.S., Consortium, F., Suzuki, H., Nefzger, C.M., Daub, C.O., et al. (2016). A predictive computational framework for direct reprogramming between human cell types. Nat Genet 48, 331–335.

Ramskold, D., Luo, S., Wang, Y.C., Li, R., Deng, Q., Faridani, O.R., Daniels, G.A., Khrebtukova, I., Loring, J.F., Laurent, L.C., et al. (2012). Full-length mRNA-Seq from single-cell levels of RNA and individual circulating tumor cells. Nat Biotechnol 30, 777–782.

Robinson, M.D., Mccarthy, D.J., and Smyth, G.K. (2010). edgeR: a Bioconductor package for differential expression analysis of digital gene expression data. 26, 139–140.

Ryoji Amamoto, and Arlotta, P. (2014). Development-Inspired Reprogramming of the Mammalian Central Nervous System. Science (80-.). 343.

Satija, R., Farrell, J.A., Gennert, D., Schier, A.F., and Regev, A. (2015). Spatial reconstruction of single-cell gene expression data. Nat. Biotechnol. 33, 495–502.

Schuurmans, C., and Guillemot, F. (2002). Molecular mechanisms underlying cell fate specification in the developing telencephalon. Curr. Opin. Neurobiol. 12, 26–34.

Shalem, O., Sanjana, N.E., and Zhang, F. (2015). High-throughput functional genomics using CRISPR-Cas9. Nat. Rev. Genet. 16, 299–311.

Shannon, P., Markiel, A., Ozier, O., Baliga, N.S., Wang, J.T., Ramage, D., Amin, N., Schwikowski, B., and Ideker, T. (2003). Cytoscape: a software environment for integrated models of biomolecular interaction networks. Genome Res 13, 2498–2504.

Shin, J.W., Suzuki, T., Ninomiya, N., Kishima, M., Hasegawa, Y., Kubosaki, A., Yabukami, H., Hayashizaki, Y., and Suzuki, H. (2012). Establishment of single-cell screening system for the rapid identification of transcriptional modulators involved in direct cell reprogramming. Nucleic Acids Res 40, e165.

Smith, D.K., Yang, J., Liu, M.L., and Zhang, C.L. (2016). Small Molecules Modulate Chromatin Accessibility to Promote NEUROG2-Mediated Fibroblast-to-Neuron Reprogramming. Stem Cell Reports 7, 955–969.

Son, E.Y., Ichida, J.K., Wainger, B.J., Toma, J.S., Rafuse, V.F., Woolf, C.J., and Eggan, K. (2011). Conversion of mouse and human fibroblasts into functional spinal motor neurons. Cell Stem Cell 9, 205–218.

Stoykova, A., Treichel, D., Hallonet, M., and Gruss, P. (2000). Pax6 modulates the dorsoventral patterning of the mammalian telencephalon. J Neurosci 20, 8042–8050.

Thoma, E.C., Wischmeyer, E., Offen, N., Maurus, K., Sirén, A.L., Schartl, M., and Wagner, T.U. (2012). Ectopic expression of neurogenin 2 alone is sufficient to induce differentiation of embryonic stem cells into mature neurons. PLoS One 7.

Trapnell, C., Cacchiarelli, D., Grimsby, J., Pokharel, P., Li, S., Morse, M., Lennon, N.J., Livak, K.J., Mikkelsen, T.S., and Rinn, J.L. (2014). The dynamics and regulators of cell fate decisions are revealed by pseudotemporal ordering of single cells. Nat Biotechnol 32, 381–386.

Treutlein, B., Lee, Q.Y., Camp, J.G., Mall, M., Koh, W., Shariati, S.A., Sim, S., Neff, N.F., Skotheim, J.M., Wernig, M., et al. (2016). Dissecting direct reprogramming from fibroblast to neuron using single-cell RNA-seq. Nature 534, 391–395.

Tsunemoto, R., Lee, S., Szűcs, A., Chubukov, P., Sokolova, I., Blanchard, J.W., Eade, K.T., Bruggemann, J., Wu, C., Torkamani, A., et al. (2018). Diverse reprogramming codes for neuronal identity. Nature.

Victor, M.B., Richner, M., Hermanstyne, T.O., Ransdell, J.L., Sobieski, C., Deng, P.Y., Klyachko, V.A., Nerbonne, J.M., and Yoo, A.S. (2014). Generation of human striatal neurons by microRNA-dependent direct conversion of fibroblasts. Neuron 84, 311–323.

Vierbuchen, T., and Wernig, M. (2011). Direct lineage conversions: unnatural but useful? Nat Biotechnol 29, 892–907.

Vierbuchen, T., Ostermeier, A., Pang, Z.P., Kokubu, Y., Südhof, T.C., and Wernig, M. (2010). Direct conversion of fibroblasts to functional neurons by defined factors. Nature 463, 1035–1041.

Wapinski, O.L., Vierbuchen, T., Qu, K., Lee, Q.Y., Chanda, S., Fuentes, D.R., Giresi, P.G., Ng, Y.H., Marro, S., Neff, N.F., et al. (2013). XHierarchical mechanisms for direct reprogramming of fibroblasts to neurons. Cell 155, 621–635.

Wickham, H. (2007). Reshaping Data with the reshape Package. 21.

Xu, Z., Jiang, H., Zhong, P., Yan, Z., Chen, S., and Feng, J. (2016). Direct conversion of human fibroblasts to induced serotonergic neurons. Mol. Psychiatry 21, 62–70.

Xue, Y., Ouyang, K., Huang, J., Zhou, Y., Ouyang, H., Li, H., Wang, G., Wu, Q., Wei, C., Bi, Y., et al. (2013). Direct conversion of fibroblasts to neurons by reprogramming PTB-regulated microRNA circuits. Cell 152, 82–96.

Yan, Y., Yang, D., Zarnowska, E.D., Du, Z., Werbel, B., Valliere, C., Pearce, R.A., Thomson, J.A., and Zhang, S.C. (2005). Directed differentiation of dopaminergic neuronal subtypes from human embryonic stem cells. Stem Cells 23, 781–790.

Yun, K., Potter, S., and Rubenstein, J.L. (2001). Gsh2 and Pax6 play complementary roles in dorsoventral patterning of the mammalian telencephalon. Development 128, 193–205.

Zhang, X., Huang, C.T., Chen, J., Pankratz, M.T., Xi, J., Li, J., Yang, Y., Lavaute, T.M., Li, X.J., Ayala, M., et al. (2010). Pax6 is a human neuroectoderm cell fate determinant. Cell Stem Cell 7, 90–100.

Zheng, G.X., Terry, J.M., Belgrader, P., Ryvkin, P., Bent, Z.W., Wilson, R., Ziraldo, S.B., Wheeler, T.D., McDermott, G.P., Zhu, J., et al. (2017). Massively parallel digital transcriptional profiling of single cells. Nat Commun 8, 14049.

