## Supplementary Data for "Decoding neuronal diversity by single-cell Convert-seq"

### FIGURE S1

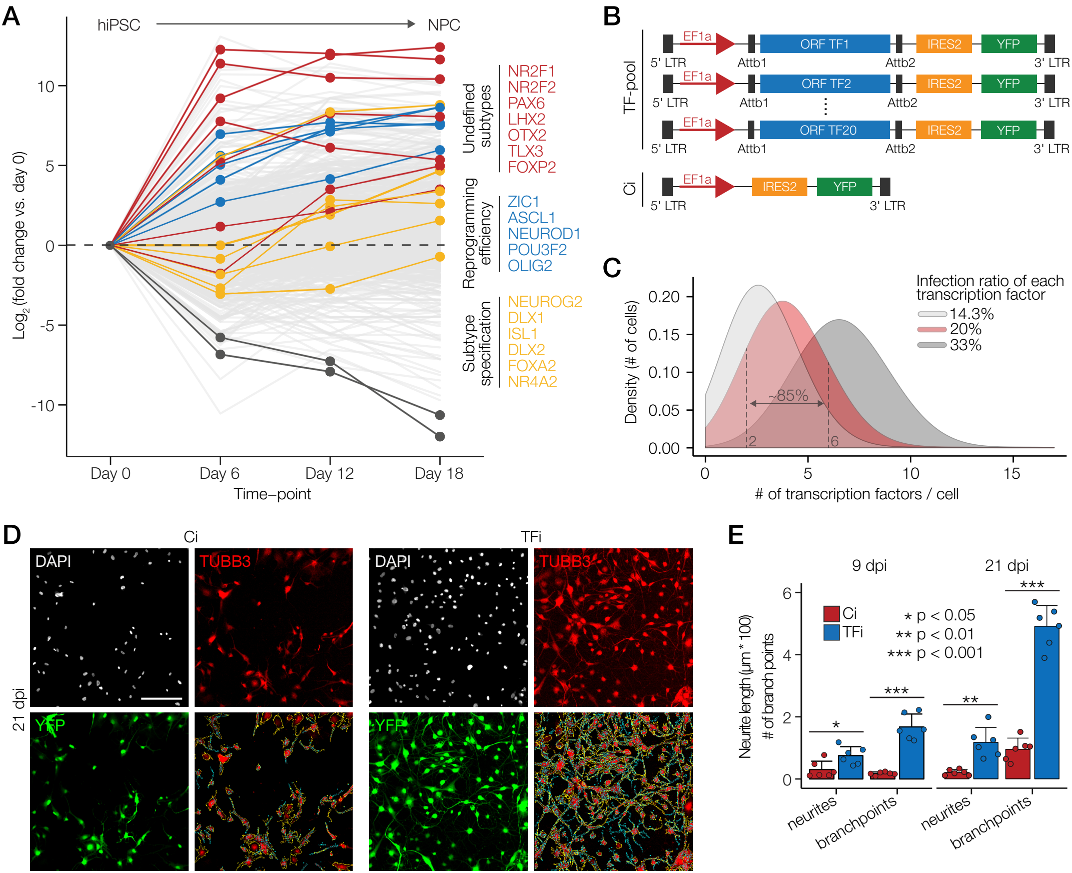
A
hiPSC
NPC
Log2 (fold change vs. day 0)
10
5
0
-5
-10
Day 0
Day 6
Day 12
Day 18
Time–point
Undefined subtypes
NR2F1
NR2F2
PAX6
LHX2
OTX2
TLX3
FOXP2
Reprogramming efficiency
ZIC1
ASCL1
NEUROD1
POU3F2
OLIG2
Subtype specification
NEUROG2
DLX1
ISL1
DLX2
FOXA2
NR4A2
B
TF-pool
EF1a
ORF TF1
IRES2
YFP
5' LTR
Attb1
Attb2
3' LTR
ORF TF2
ORF TF20
Ci
C
Density (# of cells)
0.20
0.15
0.10
0.05
0.00
Infection ratio of each transcription factor
14.3%
20%
33%
~85%
2
6
0
5
10
15
### of transcription factors / cell
D
Ci
TFi
DAPI
TUBB3
YFP
21 dpi
E
9 dpi
21 dpi
Neurite length (µm * 100)
### of branch points
6
4
2
0
Ci
TFi
* p < 0.05
** p < 0.01
*** p < 0.001
*
***
**
***
neurites
branchpoints
neurites
branchpoints

### FIGURE S2

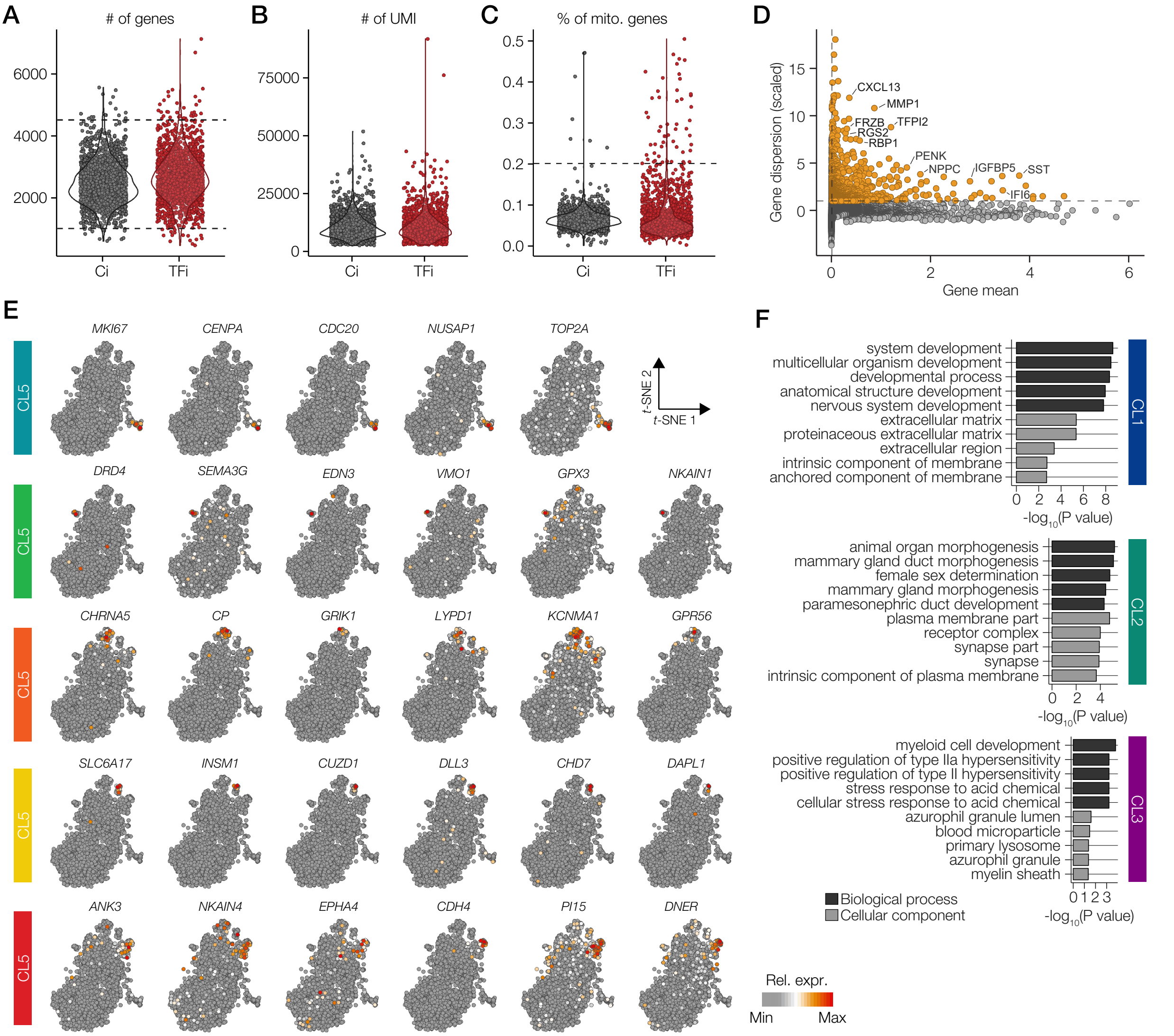

A
### of genes
6000
4000
2000
Ci
TFi
B
### of UMI
75000
50000
25000
0
Ci
TFi
C
% of mito. genes
0.5
0.4
0.3
0.2
0.1
0.0
Ci
TFi
D
Gene dispersion (scaled)
15
10
5
0
CXCL13
MMP1
FRZB
TFPI2
RGS2
RBP1
PENK
NPPC
IGFBP5
SST
IFI6
0
2
4
6
Gene mean
E
MKI67
CENPA
CDC20
NUSAP1
TOP2A
t-SNE 2
t-SNE 1
DRD4
SEMA3G
EDN3
VMO1
GPX3
NKAIN1
CHRNA5
CP
GRIK1
LYPD1
KCNMA1
GPR56
SLC6A17
INSM1
CUZD1
DLL3
CHD7
DAPL1
ANK3
NKAIN4
EPHA4
CDH4
PI15
DNER
CL5
Rel. expr.
Min
Max
F
system development
multicellular organism development
developmental process
anatomical structure development
nervous system development
extracellular matrix
proteinaceous extracellular matrix
extracellular region
intrinsic component of membrane
anchored component of membrane
0 2 4 6 8
-log10(P value)
CL1
animal organ morphogenesis
mammary gland duct morphogenesis
female sex determination
mammary gland morphogenesis
paramesonephric duct development
plasma membrane part
receptor complex
synapse part
synapse
intrinsic component of plasma membrane
0 2 4
-log10(P value)
CL2
myeloid cell development
positive regulation of type IIa hypersensitivity
positive regulation of type II hypersensitivity
stress response to acid chemical
cellular stress response to acid chemical
azurophil granule lumen
blood microparticle
primary lysosome
azurophil granule
myelin sheath
0 1 2 3
-log10(P value)
CL3
Biological process
Cellular component

FIGURE S3

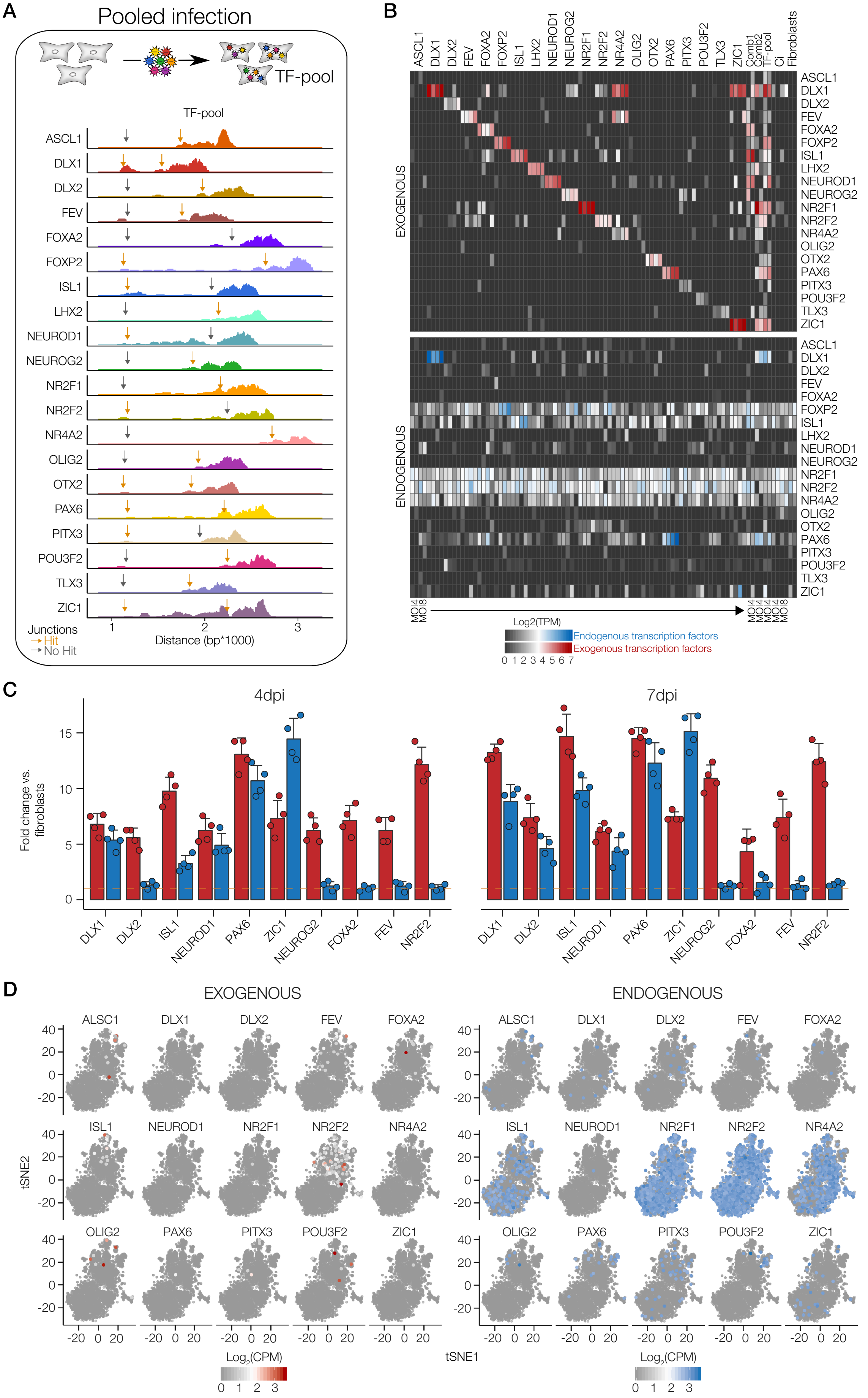
A
Pooled infection
TF-pool
TF-pool
ASCL1
DLX1
DLX2
FEV
FOXA2
FOXP2
ISL1
LHX2
NEUROD1
NEUROG2
NR2F1
NR2F2
NR4A2
OLIG2
OTX2
PAX6
PITX3
POU3F2
TLX3
ZIC1
Junctions
Hit
No Hit
1
2
3
Distance (bp*1000)
B
ASCL1 DLX1 DLX2 FEV FOXA2 FOXP2 ISL1 LHX2 NEUROD1 NEUROG2 NR2F1 NR2F2 NR4A2 OLIG2 OTX2 PAX6 PITX3 POU3F2 TLX3 ZIC1 Comb1 Comb2 TF-pool Ci Fibroblasts
EXOGENOUS
ENDOGENOUS
MOI4
MOI8
Log2(TPM)
Endogenous transcription factors
Exogenous transcription factors
0 1 2 3 4 5 6 7
C
4dpi
7dpi
Fold change vs. fibroblasts
0
5
10
15
DLX1
DLX2
ISL1
NEUROD1
PAX6
ZIC1
NEUROG2
FOXA2
FEV
NR2F2
D
EXOGENOUS
ENDOGENOUS
ALSC1
DLX1
DLX2
FEV
FOXA2
ISL1
NEUROD1
NR2F1
NR2F2
NR4A2
OLIG2
PAX6
PITX3
POU3F2
ZIC1
tSNE2
tSNE1
40
20
0
-20
-20 0 20
Log2(CPM)
0 1 2 3

#### FIGURE S4

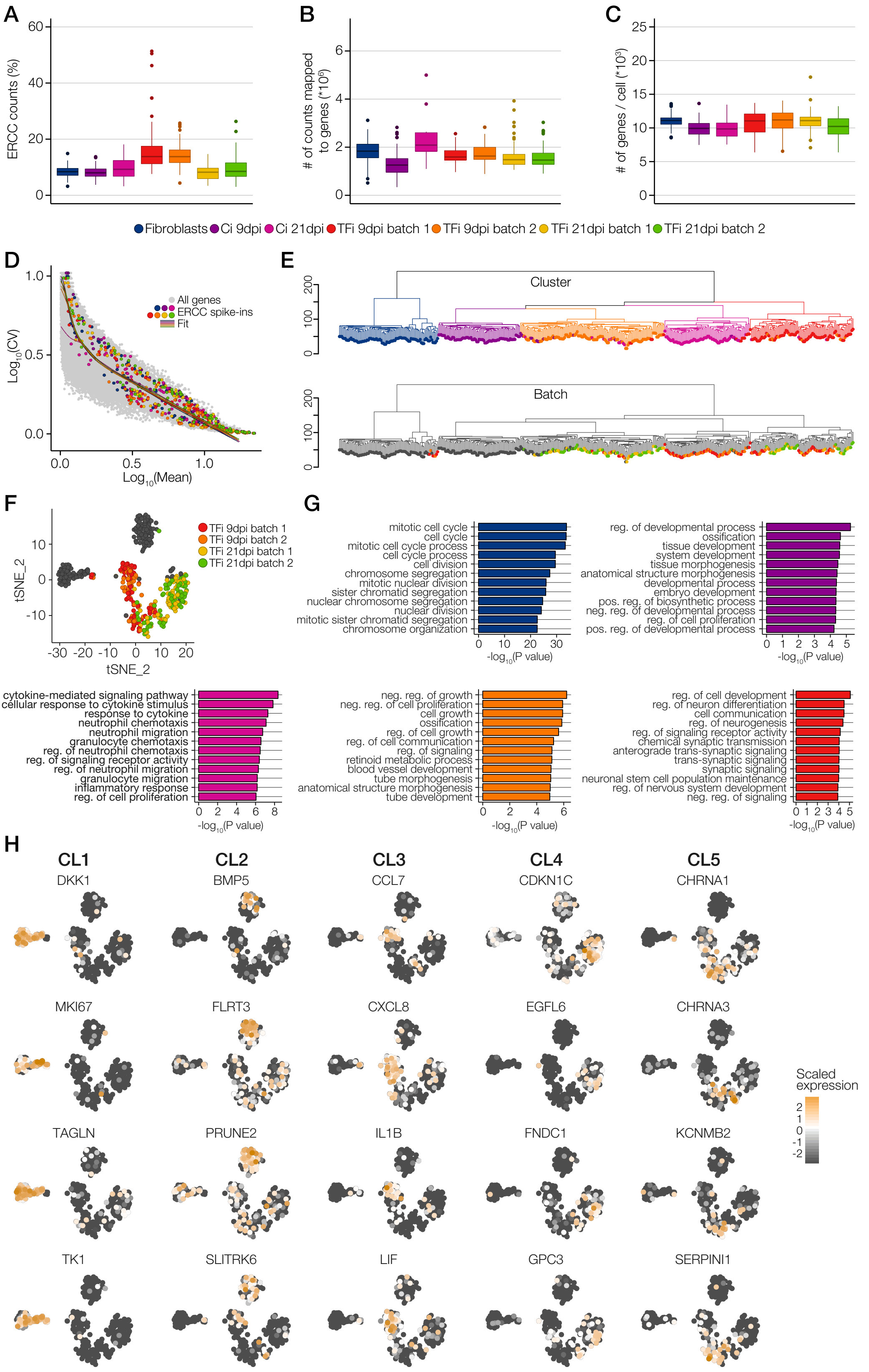
A
ERCC counts (%)
B
### of counts mapped to genes (*10^6)
C
### of genes / cell (*10^3)
Fibroblasts
Ci 9dpi
Ci 21dpi
TFi 9dpi batch 1
TFi 9dpi batch 2
TFi 21dpi batch 1
TFi 21dpi batch 2
D
All genes
ERCC spike-ins
Fit
Log10(CV)
Log10(Mean)
E
Cluster
Batch
F
TFi 9dpi batch 1
TFi 9dpi batch 2
TFi 21dpi batch 1
TFi 21dpi batch 2
tSNE_2
tSNE_2
G
mitotic cell cycle
cell cycle
mitotic cell cycle process
cell cycle process
cell division
chromosome segregation
mitotic nuclear division
sister chromatid segregation
nuclear chromosome segregation
nuclear division
mitotic sister chromatid segregation
chromosome organization
-log10(P value)
reg. of developmental process
ossification
tissue development
system development
tissue morphogenesis
anatomical structure morphogenesis
developmental process
embryo development
pos. reg. of biosynthetic process
neg. reg. of developmental process
reg. of cell proliferation
pos. reg. of developmental process
-log10(P value)
cytokine-mediated signaling pathway
cellular response to cytokine stimulus
response to cytokine
neutrophil chemotaxis
neutrophil migration
granulocyte chemotaxis
reg. of neutrophil chemotaxis
reg. of signaling receptor activity
reg. of neutrophil migration
granulocyte migration
inflammatory response
reg. of cell proliferation
-log10(P value)
neg. reg. of growth
neg. reg. of cell proliferation
cell growth
ossification
reg. of cell growth
reg. of cell communication
reg. of signaling
retinoid metabolic process
blood vessel development
tube morphogenesis
anatomical structure morphogenesis
tube development
-log10(P value)
reg. of cell development
reg. of neuron differentiation
cell communication
reg. of neurogenesis
reg. of signaling receptor activity
chemical synaptic transmission
anterograde trans-synaptic signaling
trans-synaptic signaling
synaptic signaling
neuronal stem cell population maintenance
reg. of nervous system development
neg. reg. of signaling
-log10(P value)
H
CL1
CL2
CL3
CL4
CL5
DKK1
BMP5
CCL7
CDKN1C
CHRNA1
MKI67
FLRT3
CXCL8
EGFL6
CHRNA3
TAGLN
PRUNE2
IL1B
FNDC1
KCNMB2
TK1
SLITRK6
LIF
GPC3
SERPINI1
Scaled expression

### FIGURE S5

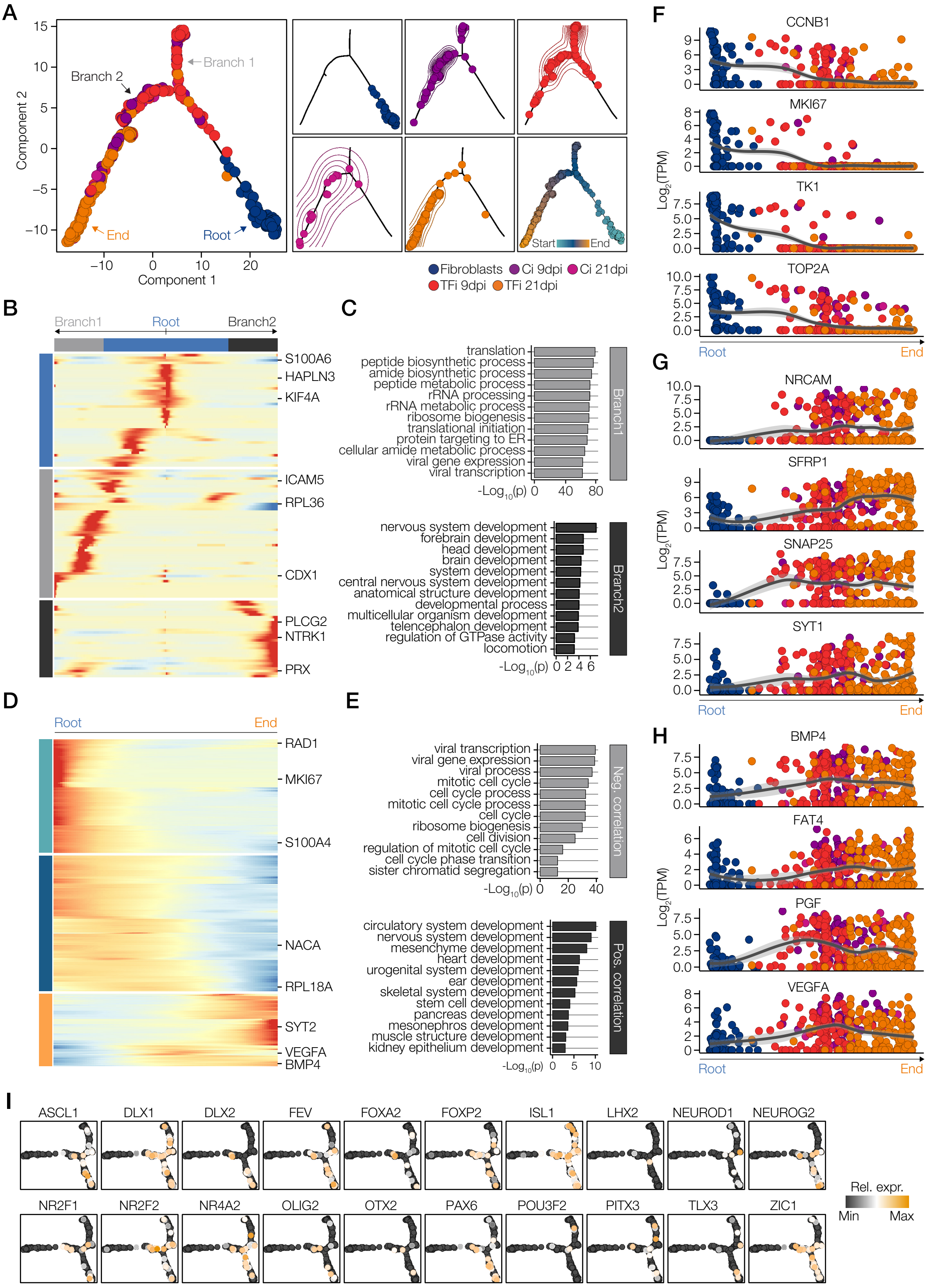
A
Component 2
Component 1
Branch 1
Branch 2
End
Root
Start
End
Fibroblasts
Ci 9dpi
Ci 21dpi
TFi 9dpi
TFi 21dpi
B
Branch1
Root
Branch2
S100A6
HAPLN3
KIF4A
ICAM5
RPL36
CDX1
PLCG2
NTRK1
PRX
C
translation
peptide biosynthetic process
amide biosynthetic process
peptide metabolic process
rRNA processing
rRNA metabolic process
ribosome biogenesis
translational initiation
protein targeting to ER
cellular amide metabolic process
viral gene expression
viral transcription
Branch1
nervous system development
forebrain development
head development
brain development
system development
central nervous system development
anatomical structure development
developmental process
multicellular organism development
telencephalon development
regulation of GTPase activity
locomotion
Branch2
D
Root
End
RAD1
MKI67
S100A4
NACA
RPL18A
SYT2
VEGFA
BMP4
E
viral transcription
viral gene expression
viral process
mitotic cell cycle
cell cycle process
mitotic cell cycle process
cell cycle
ribosome biogenesis
cell division
regulation of mitotic cell cycle
cell cycle phase transition
sister chromatid segregation
Neg. correlation
circulatory system development
nervous system development
mesenchyme development
heart development
urogenital system development
ear development
skeletal system development
stem cell development
pancreas development
mesonephros development
muscle structure development
kidney epithelium development
Pos. correlation
F
CCNB1
MKI67
TK1
TOP2A
Log2(TPM)
Root
End
G
NRCAM
SFRP1
SNAP25
SYT1
H
BMP4
FAT4
PGF
VEGFA
I
ASCL1
DLX1
DLX2
FEV
FOXA2
FOXP2
ISL1
LHX2
NEUROD1
NEUROG2
NR2F1
NR2F2
NR4A2
OLIG2
OTX2
PAX6
POU3F2
PITX3
TLX3
ZIC1
Rel. expr.
Min
Max

### FIGURE S6

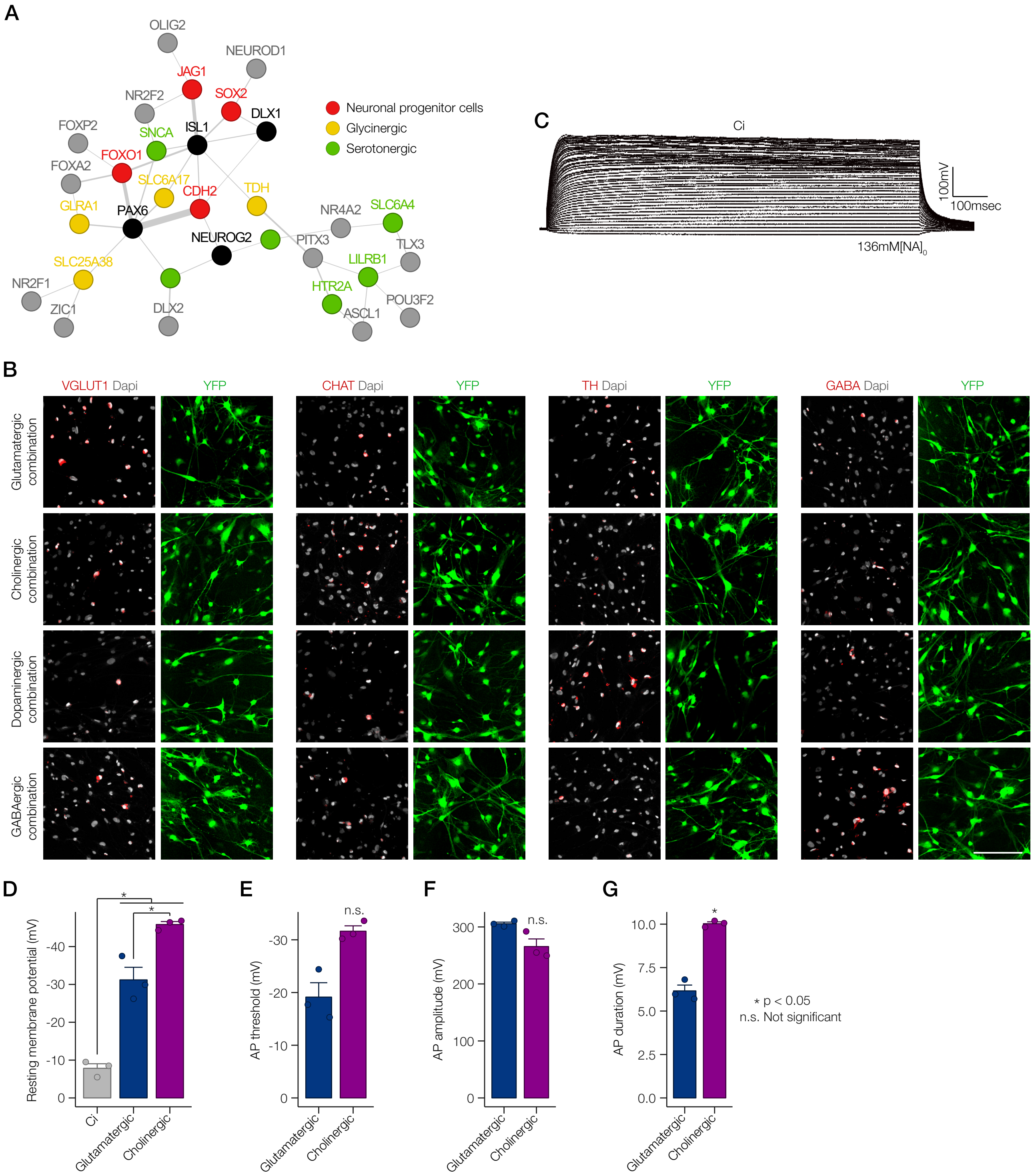

A
OLIG2
NEUROD1
JAG1
NR2F2
SOX2
DLX1
FOXP2
SNCA
ISL1
FOXO1
FOXA2
SLC6A17
CDH2
TDH
GLRA1
PAX6
NR4A2
SLC6A4
NEUROG2
PITX3
TLX3
SLC25A38
LILRB1
NR2F1
HTR2A
POU3F2
ZIC1
DLX2
ASCL1
Neuronal progenitor cells
Glycinergic
Serotonergic
C
Ci
100mV
100msec
136mM[NA]0
B
VGLUT1 Dapi
YFP
CHAT Dapi
TH Dapi
GABA Dapi
Glutamatergic combination
Cholinergic combination
Dopaminergic combination
GABAergic combination
D
Resting membrane potential (mV)
Ci
Glutamatergic
Cholinergic
E
AP threshold (mV)
n.s.
F
AP amplitude (mV)
G
AP duration (mV)
* p < 0.05
n.s. Not significant

**Figure S1, Related to Figure 1. Candidate transcription factor expression during iPSC-to-NPC differentiation and neuronal profiling of TFi and Ci.**

(A) Expression fold changes of 18 out of 20 candidate neurogenic transcription factors (colored) during differentiation of human induced pluripotent stem cells (hiPSC; day 0) into early neuronal progenitor cells (NPC; day 18). Pluripotency markers *OCT4* and *NANOG* are shown in dark grey, all other transcription factors are shown in light gray.

(B) Schematic overview of the expression vectors of the TF-pool (top) and Ci (bottom).

(C) Theoretical prediction of the number of transcription factors each cell will be infected with, assuming that each transcription factor infects 14.3% (light gray), 20% (red) and 33% (dark gray) of cells.

(D) Neuronal profiling of Ci and TFi was performed on pictures of immunostainings for TUBB3 (red).

(E) Quantification of the length of neurites and the number of branch points of Ci and TFi at 7 dpi and 21 dpi. n = 6 independent experiments, error bars represent mean + SD.

**Figure S2, Related to Figure 2. Quality control and marker gene expression of droplet-based scRNA-seq data.**

(A-C) Violin plots of the number of detected genes (A), the number of UMI (B) and the percentage of mitochondrial genes (C) of sequenced Ci (gray) and TFi (red) ($n$ = 3865 cells). Dashed lines show thresholds applied for quality control.

(D) Mean variation plot of all genes after quality control. Variable genes used for PCA and *t*-SNE are shown in red.

(E) Visualization of the cluster-specific relative expression levels of marker genes using t-SNE; cluster colors as in Figure 2A. The full list of differentially expressed genes of all clusters can be found in Supplementary Table 2.

(F) GO analysis of cluster-specific marker genes in clusters CL1 – CL3. Shown are top 5 GO terms related to biological process (dark grey) and cellular component (light grey) for each cluster.

**Figure S3, Related to Figure 3. Benchmark of bulk Convert-seq and single-cell Convert-seq of droplet-based scRNA-seq data.**

(A) Bulk Convert-seq on pooled infected fibroblasts. Horizontal dimension; distance from the 5‘ end of the EF1A promoter, vertical dimension; number of aligned paired-end reads. Gray arrows (no overlap) and golden arrows (overlap) mark 5’ and 3’ junctions of exogenous ORFs.

(B) Heat map showing $\log_2$-transformed TPM values of exogenous (red) and endogenous (blue) transcription factor pairs after alignment using Kallisto without trimming junction sequences. For individually infected fibroblasts and Ci, 2 replicates at an MOI of 4 and 2 replicates at an MOI of 8 were included. For pooled infected fibroblasts, 2 replicates at an MOI of 4 were included.

(C) Validation of up-regulation of endogenous expression of *DLX1*, *DLX2*, *ISL1*, *NEUROD1*, *PAX6* and *ZIC1* after infection of their exogenous counterparts into fibroblasts as revealed by qPCR at 4 dpi and 7 dpi using primers flanking the 3‘ junctions of ORFs. No upregulation of endogenous *NEUROG2*, *FOXA2*, *FEV* and *NR2F2* was observed. $n = 4$ independent experiments, error bars represent mean + SD.

(D) Visualization of $\log_2$-transformed CPM expression values of exogenous (red) and endogenous (blue) transcription factors on two-dimensional t-SNE projections reveal inefficient detection of exogenous transcription factors in droplet-based scRNA-seq data ($n = 3865$ cells).

**Figure S4, Related to Figure 4. Quality control and single-cell Convert-seq of Fluidigm C1 data.**

(A-C) Box plots showing the percentage of ERCC spike-ins (A), the number of counts mapped to genes (B) and the number of detected genes (C) in all 7 runs sequenced for the time-course experiment ($n = 446$ cells).

(D) Coefficient of variation is plotted against mean TPM, all genes are shown in gray, lines indicate the fit for each run and ERCC spike-ins are shown in colored dots; colors as in A.

(E) Hierarchical clustering of fibroblasts, Ci and TFi at 9 dpi and 21 dpi recapitulates 2-dimensional visualization by *t*-SNE shown in Figure 4B (top) and reveals no batch effects (bottom).

(F) *t*-SNE projection of transcriptomic data (Fluidigm C1) where TFi batches are colored and all other cells are gray. Clustering of TFi is not batch-dependent.

(G) GO analysis of cluster-specific marker genes of the time-course data. Shown are top 12 GO terms related to biological process for each cluster.

(F) Visualization of scaled expression values of selected marker genes of each cluster on two-dimensional t-SNE projections.

**Figure S5, Related to Figure 5. Unsupervised pseudo-temporal ordering of time-course data identifies parallel developmental trajectories.**

(A) Pseudo-temporal ordering of time-course data based on genes differentially expressed between 9 dpi and 21 dpi ($n$ = 446 cells). Small squares show the same plot colored by pseudo-temporal values and separate density plots for each sample.

(B) Heat map showing ~100 genes with branch-specific differential expression as determined by BEAM (R Package 'Monocle'). In this heat map, columns are points in pseudo-time, rows are genes and the middle (Root) is the beginning of pseudo-time. Branch 1 goes from the middle of the heat map to the left while branch 2 goes to the right.

(C) Top 12 GO terms enriched in genes differentially expressed in branch 1 (top panel) and branch 2 (bottom panel).

(D) Heat map showing ~1000 genes whose relative expression changes as a function of pseudo-time.

(E) Top GO terms enriched in the top 2000 genes showing negative (top panel) and positive (bottom panel) Pearson correlation with pseudo-temporal values.

(F-H) Dot plots and fit (gray) of $\log_2$-transformed TPM expression values of cell cycle-related genes (*CCNB1*, *MKI67*, *TK1*, *TOP2A*; F), canonical neuronal genes (*NRCAM*, *SFRP1*, *SNAP25*, *SYT1*; G) and genes associated with alternative developmental fates (*BMP4*, *FAT4*, *PGF*, *VEGFA*; H) along pseudo-time; colors as in A.

(I) Visualization of relative expression values of exogenous transcription factors along pseudo-time where cells are ordered based on the expression of developmental genes.

**Figure S6, Related to Figure 6. Validation of neuronal subtype-specific GRNs.**

(A) Merged GRN showing significant associations of exogenous transcription factors with NPC, glycinergic and serotonergic genes (colored). Exogenous transcription factors with at least 3 edges are shown in black, all other exogenous transcription factors are shown in gray.

(B) Representative immunostainings for VGLUT1, CHAT, TH and GABA of fibroblasts infected with exogenous transcription factors which associated with glutamatergic, cholinergic, dopaminergic and GABAergic GRNs. Scale bar, 100μm. YFP (green) marks infected cells and cell nuclei were visualized using DAPI nuclear stain (grey).

(C) The generation of the action potential in control cells (Ci). Representative traces in the presence of extracellular $Na^+$ were recorded using the current-clamp protocol.

(D-G) Electrophysiological properties of control cells (Ci), iN infected with *DLX2*, *NEUROG2*, *PAX6*, *ZIC1* (glutamatergic) or *DLX1*, *ISL1*, *NEUROG2*, *PAX6* (cholinergic).
